# Comprehensive single-cell genome analysis at nucleotide resolution using the PTA Analysis Toolbox

**DOI:** 10.1101/2023.02.15.528636

**Authors:** Sjors Middelkamp, Freek Manders, Flavia Peci, Markus J. van Roosmalen, Diego Montiel González, Eline J.M. Bertrums, Inge van der Werf, Lucca L.M. Derks, Niels M. Groenen, Mark Verheul, Laurianne Trabut, Arianne M. Brandsma, Evangelia Antoniou, Dirk Reinhardt, Marc Bierings, Mirjam E. Belderbos, Ruben van Boxtel

## Abstract

Detection of somatic mutations in single cells has been severely hampered by technical limitations of whole genome amplification. Novel technologies including primary template-directed amplification (PTA) significantly improved the accuracy of single-cell whole genome sequencing (WGS), but still generate hundreds of artefacts per amplification reaction. We developed a comprehensive bioinformatic workflow, called the PTA Analysis Toolkit (PTATO), to accurately detect single base substitutions, small insertions and deletions (indels) and structural variants in PTA-based WGS data. PTATO includes a machine learning approach to distinguish PTA-artefacts from true mutations with high sensitivity (up to 90% for base substitution and 95% for indels), outperforming existing bioinformatic approaches. Using PTATO, we demonstrate that many hematopoietic stem and progenitor cells of patients with Fanconi anemia, which cannot be analyzed using regular WGS technologies, have normal somatic single base substitution burdens, but increased numbers of deletions. Our results show that PTATO enables studying somatic mutagenesis in the genomes of single cells with unprecedented sensitivity and accuracy.

## Introduction

Somatic mutations gradually accumulate in each cell during life, which can contribute to the development of age-related diseases, such as cancer^1–3^. Due to the stochastic nature of mutation accumulation, each cell contains a unique set of somatic variants. Amplification of the genome of a single cell is required to obtain sufficient DNA for WGS. One approach for this is to catalogue mutations in clonal structures that exist in tissues *in vivo*^4^ or after clonally expanding single cells isolated from tissues *in vitro*^5,6^. However, these approaches can only be applied to cells that have the capacity to clonally expand such as stem cells, precluding analyses of many diseased and/or post-mitotic differentiated cell types^7^. Examples of these are hematopoietic stem and progenitor cells (HSPCs) of patients with Fanconi anemia (FA), who suffer from progressive bone marrow failure and are predisposed to cancer due to an inherited deficiency of DNA repair^8–10^. Much of the research into the mutagenic processes in FA HSPCs has been performed using mouse models^11–13^, because primary HSPCs of human patients with FA are difficult to culture and clonally expand *in vitro*^14,15^.

An alternative method to clonal expansion is the use of whole genome amplification (WGA) techniques to directly amplify DNA of single cells in enzymatic reactions. However, single-cell WGA technologies have traditionally been hindered by technical limitations due to uneven and erroneous amplification of the genome, leading to artificial mutations, noise in copy number profiles and missing mutations due to allelic dropout^16^. Recently, a novel WGA method, called primary template-directed amplification (PTA), was developed, which contains several critical improvements over the traditionally used multiple displacement amplification (MDA) protocol^17^. Although the amplification biases and allelic dropout rates of PTA are remarkably low, it still generates hundreds to thousands of false positive single base substitutions and indels in each amplification reaction^17,18^. Bioinformatic approaches, such as linked-read analysis (LiRA) and SCAN2, have been developed to filter and analyze WGS data of WGA samples^18,19^. However, these tools still have low detection sensitivities (~10-40%) and therefore most true variants are missed^18,19^. Additionally, while PTA has the potential to enable structural variant (SV) detection in single cells, current tools are not optimized for PTA-based single-cell WGS data.

Here, we developed the PTA Analysis Toolbox (PTATO), which uses a machine learning model to accurately filter artefacts from PTA-based WGS data and is optimized for SV detection. We demonstrate the applicability of PTATO by analyzing the genomes of normal HSPCs of FA patients and show that, similar to current FA mouse models, these cells have an increased somatic deletion burden.

## Results

### Training a random forest model to filter PTA artefacts

The artefacts generated by PTA have been shown to follow a specific, non-random 96-trinucleotide mutational profile in WGS data^17,18^. We hypothesized that we could use a machine learning approach to distinguish PTA artefacts from true positive variants based on multiple genomic features (Figure 1a). For this, we trained a random forest (RF) model, which we previously showed to be highly effective in attributing individual mutations to a specific mutational process^20^. To generate a confident set of true positive somatic variants for training of the classifier, we sequenced samples of patients with acute myeloid leukemia (AML) and cell lines using regular bulk WGS as well as single-cell WGS after PTA (Figure 1b and Supplementary Table S1). Somatic variants that were shared between the bulk and single-cell sequenced samples were used as high confidence true variants. To obtain a high confidence set of PTA artefacts for training, we used PTA-based WGS of umbilical cord blood-derived HSPCs. Most of the unique somatic variants in these cells will be PTA artefacts, because HSPCs at birth only harbor 20-50 somatic mutations^21–23^. In total, 756 PTA artefacts and 756 true positive single base substitutions were used to train the random forest model (Figure 1b). To train the RF model, we used a variety of genomic features, such as the level of allelic imbalance of the region the variant is located in, the 10-basepair (bp) sequence context around the variant, the distance to the nearest gene and replication timing (Figure 1c).

**Figure 1.**
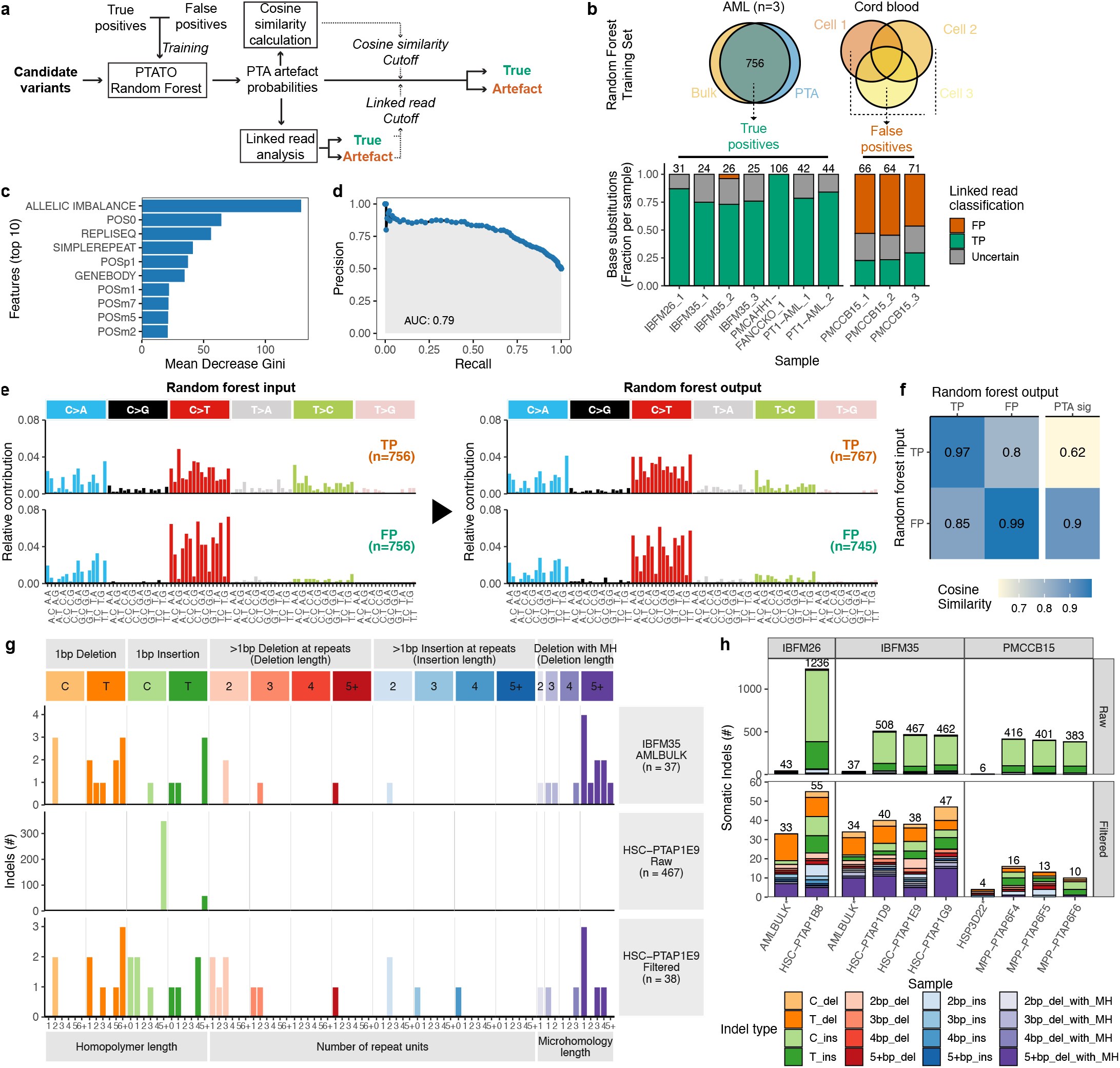
Accurate filtering of PTA artefacts using machine learning and recurrency filtering. **a,** Outline of the PTATO workflow to classify candidate base substitutions as true variants or PTA artefacts. The trained PTATO RF model calculates the probability that each variant is a PTA artefact. Subsequently it uses a linked read analysis and cosine similarity calculations to determine a sample-specific probability cutoff. **b,** Overview of the samples and variants (with their linked read classifications) that are used as PTA artefacts or true variants to train the RF model. Numbers above the bars indicate the number of variants per sample that could be analyzed by the linked read approach. **c,** Importance of the top 10 features used by the RF model to distinguish true variants from PTA artefacts. POS indicates the genomic position in base pairs relative to the mutation (m = minus, p = plus). **d,** Precision-recall curve showing the performance of the random forest using all input variables on the out-of-bag training data for different probability cutoffs. **e,** The 96-trinucleotide mutational spectra of the base substitutions that were used as PTA artefact or true positive input for training the RF model (left) and the profiles of the base substitutions that were classified as true or false by the model during cross-validation (right). **f,** Heatmap showing the cosine similarities between the base substitutions used in the training set and the base substitutions classified during cross-validation and the previously defined mutational signature of PTA artefacts. **g,** Spectra of indels detected in bulk WGS data of AML blasts (top) or before (center) and after (bottom) PTATO filtering of PTA-based WGS data of a HSPC of the same individual. **h,** Numbers and types of indels detected before (top) and after (bottom) PTATO filtering. MH, microhomology.

The RF model calculates a probability score that a candidate variant is a PTA artefact. As the PTA efficiency can vary between samples^17^, a sample-specific cutoff needs to be set above which variants are classified as artefacts. To set an optimal cut-off for each sample, we applied two complementary methods (Figure 1a and Figure S1a-e). First, PTATO uses a linked read analysis, which is a method to detect artefacts with high specificity, but low sensitivity^19^, to classify the small subset of somatic variants that can be linked to informative germline variants as true or false positive. Next, it takes the PTA probability scores for all the variants classified by the linked read analysis and calculates precision-recall curves to determine the optimal cutoff to discriminate these two groups (Figure S1c,d). Although this method works well to determine an optimal PTA probability cutoff for most samples, we noted that for some samples accurate precision-recall curves could not be generated because these samples have too few informative true variants (Figure S1c,d). Therefore, we included a second method which calculates mutational spectra at varying PTA probabilities and determines the cosine similarities between these spectra. Subsequently, the cutoff is calculated by hierarchical clustering to separate one cluster with similar mutational spectra and low probability scores (containing true variants) from a cluster of high probability scores (containing artefacts) (Figure S1e). The RF model was predicted to distinguish artefacts from true positive variants relatively well with an out-of-bag error rate of 0.264 and an area-under-the-curve for precision-recall rates of 0.79 for single base substitutions (Figure 1d and Figure S1f). Importantly, the 96-trinucleotide mutational spectra of the variants predicted to be false or true variants were nearly the same as the profiles of the input PTA artefacts or true positive variants, respectively (Figure 1e,f).

Compared to the base substitution artefacts, the indel artefacts caused by PTA follow an even more specific pattern, which is mainly characterized by C- or T-insertions at long homopolymers (repeats of the same nucleotide) (Figure 1g,h)^18^. We found that exclusively filtering indel artefacts that are recurrently called in multiple unrelated individuals and filtering insertions at long (5bp+) homopolymers was even more effective than training a RF model for indel filtering (Figure S2a,b). Indeed, this former approach removed most indel artefacts, leading to indel burdens and patterns that were comparable between those found in bulk and PTA-based WGS data (Figure 1g,h). Thus, these initial validations demonstrate that PTATO can accurately discriminate true and false positive base substitutions as well as indels using machine learning classification and filtering based on recurrence, respectively.

### Validation of the random forest model

To further test the performance of PTATO on samples that were not used in the training set, we inactivated the *FANCC* and *MSH2* genes in the human AHH-1 lymphoblast cell line using CRISPR/Cas9 gene editing (Figure S3a-c). Inactivation of these genes and their associated DNA repair pathways has been shown to induce various specific base substitution and indel signatures^24–26^, enabling us to test the performance of PTATO on a variety of mutational outcomes. We performed several sequential clonal steps followed by regular bulk and single-cell PTA-based WGS to characterize the *in vitro* accumulation of mutations in these cells (Figure 2a and Supplementary Table S2). Bulk WGS of the subclones was used as a control, which showed that the wildtype, *FANCC*^-/-^ and *MSH2*^-/-^ AHH1 clones acquire respectively 10.6, 10.5 and 52.6 base substitutions and 1.02, 1.12 and 91.1 indels per day in culture on average (Figure S4a and Figure 5a). The standard somatic variant calling pipeline (Methods) without PTATO filtering detected a 1.37-1.86 fold higher base substitution rate and a 12-29 fold higher indel rate in the PTA-amplified wildtype and *FANCC*^-/-^ samples compared to the subclones analyzed by bulk WGS (Figure 2b,c, Figure S4b and Figure 5b-c). PTATO removed most excess mutations, leading to similar mutation rates between the bulk WGS-analyzed subclones and the PTA samples (Figure 2b,c, Figure S4a-b and Figure S5a-c). Filtering by PTATO also improved the similarity between the mutational profiles of the PTA-amplified samples and the profiles of the corresponding bulk WGS-analyzed subclones (Figure 2d, Figure S4c-g and Figure S5d-e). As shown for the *MSH2*^-/-^ cell sequenced after PTA, PTATO can also accurately remove PTA artefacts from samples with low amplification quality, although the sensitivity to detect true variants is reduced due to uneven coverage of the genome (Figure 2b-g, Figure S4 and Figure S5).

**Figure 2.**
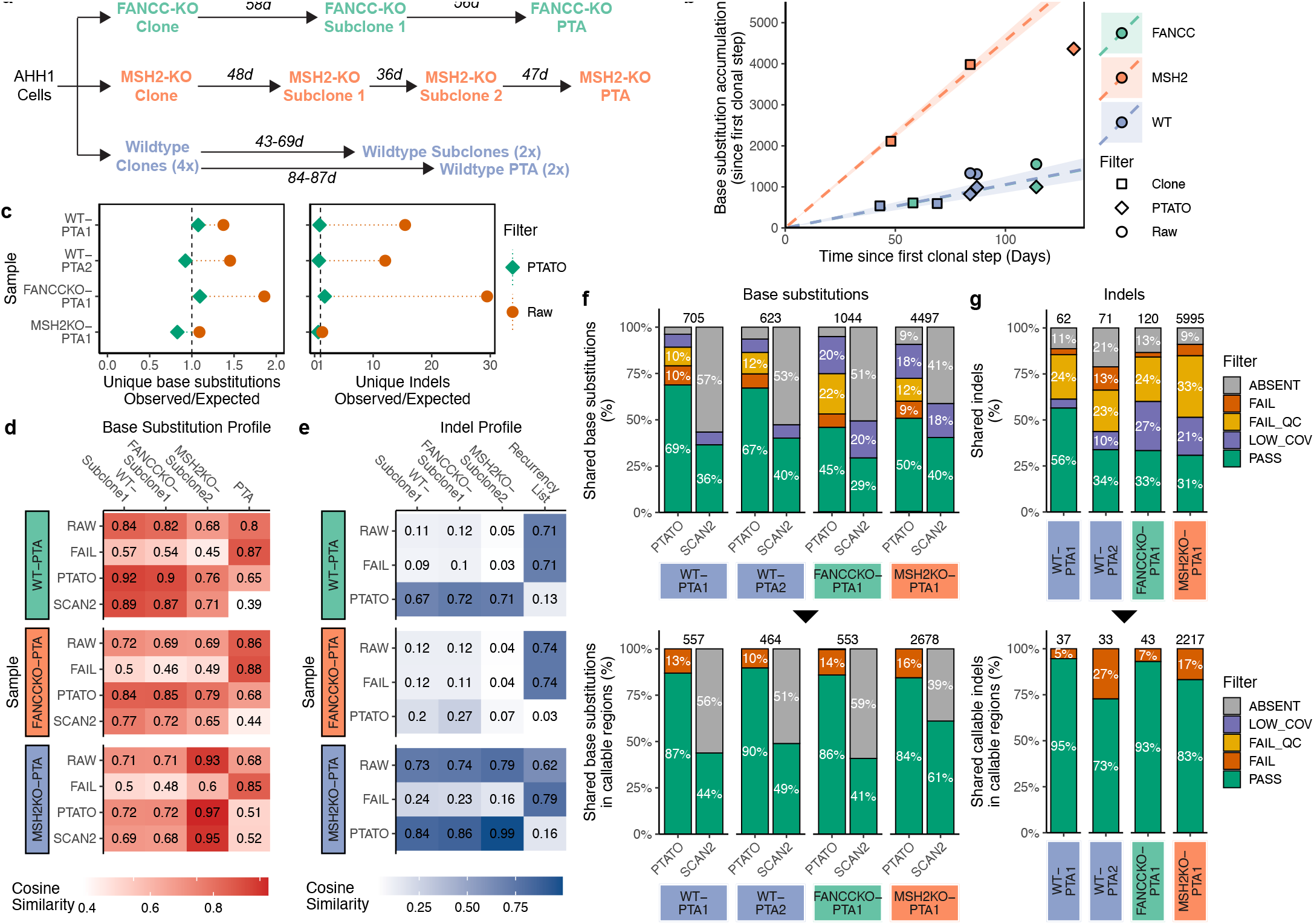
Filtering by PTATO enables accurate analyses of somatic mutation patterns and burdens. **a,** Schematic overview of the clonal steps performed for the three types of clonal cell lines generated in this study. Numbers indicate the days (d) in culture between the single cell sorts, which are used to calculate mutation rates for each cell line. **b,** Accumulation of base substitutions per sample since the first clonal step. The circles and diamonds indicate the number of base substitutions detected in the PTA samples before and after PTATO filtering, respectively. **c,** Observed versus expected number of base substitutions (left) and indels (right) in the PTA samples before (orange) and after (green) filtering by PTATO. **d,** Heatmap showing the cosine similarities between the 96-trinucleotide profiles of the unique base substitutions before PTATO filtering (RAW), after PTATO filtering, after SCAN2 calling or the mutations removed by PTATO (FAIL) and the profiles of the subclones analyzed by bulk WGS or the previously defined PTA artefact signature. **e,** Heatmap showing the cosine similarities between the profiles of the unique indels before PTATO filtering (RAW), after PTATO filtering or the mutations removed by PTATO (FAIL) and the indel profiles of the subclones analyzed by bulk WGS or the list of recurrent indels used for filtering. **f-g,** Fractions of base substitutions (f) and indels (g) present in the subclones that are also detected (PASS) in the PTA samples originating from these subclones by PTATO or SCAN2 (SCAN2 could not be used to study indels in these samples). Bottom panels show the base substitutions (f) and indels (g) after excluding the variants with low coverage (LOW_COV), low genotype quality (LOW_QC) or undetected variants (ABSENT). Few shared variants are (mis)classified as artefact (FAIL) in the PTA samples.

**Figure 3.**
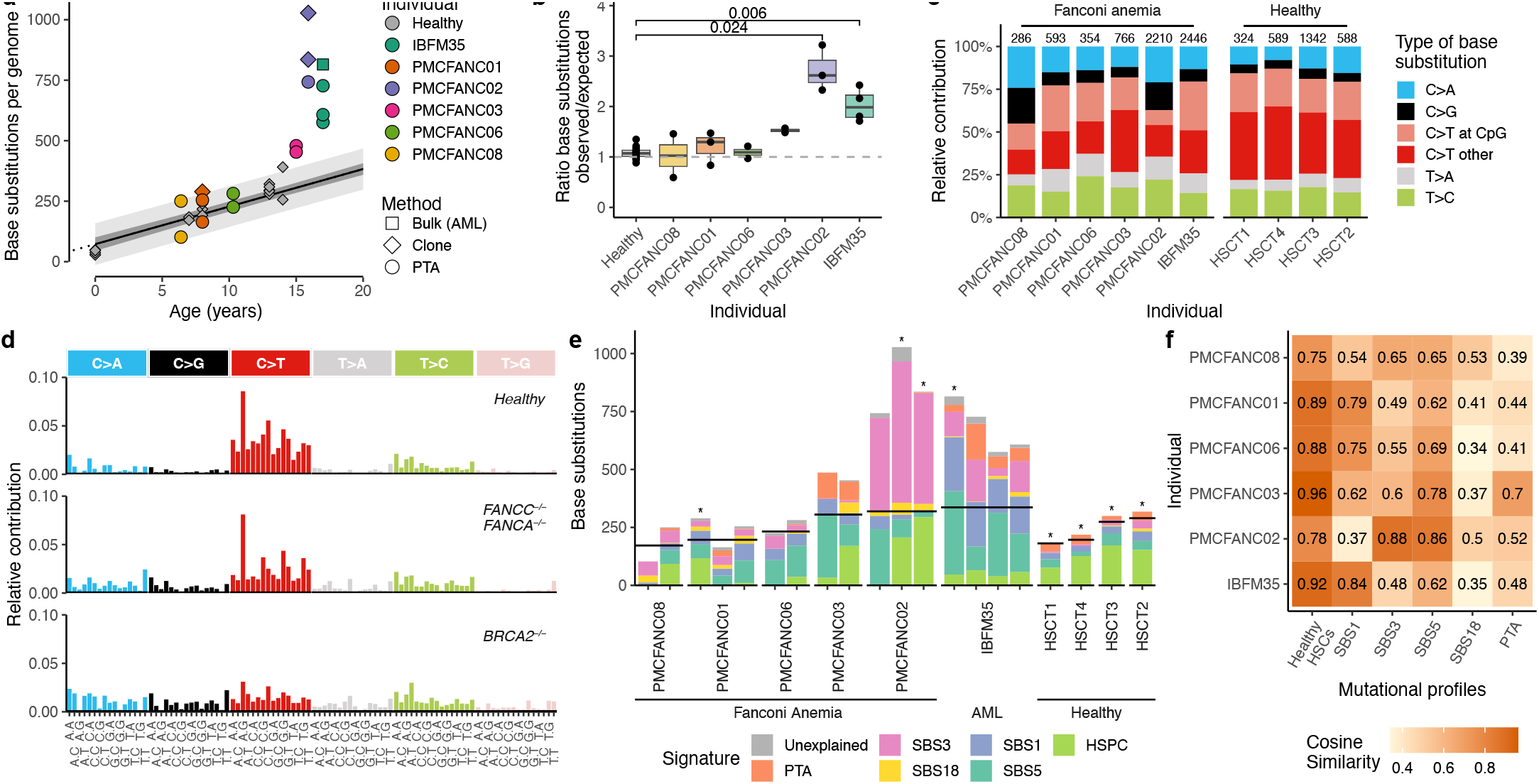
PTATO detects normal single base substitution burdens in most human FA HSPCs. **a,** Correlation of the number of single base substitutions per HSPC genome of healthy donors (gray points) and patients with FA. Linear mixed modelling showed that healthy HSPCs accumulate base substitutions in a linear fashion with age^21,22^. The 95% confidence interval and the prediction interval of the model are indicated by the dark gray and light gray shading, respectively. **b,** Ratios between the observed and expected number of base substitutions per genome (sorted on age) based on extrapolation of the age linear mixed model. To match the ages of the patients with FA, only 12 HSPCs of four healthy donors (HSCT1-4, ages 7 to 14) are included in this and following panels. Adjusted P-values indicate multiple testing corrected significant differences (padj<0.05) between three FA patients and the age-matched healthy donors (Bonferroni-corrected Wilcoxon Mann–Whitney test). **c,** Mutation spectra showing the relative contribution of each base substitution type in the genomes of the donors. Numbers above the bar indicate the total number of base substitutions found in the samples from each individual. **d,** The averaged 96-trinucleotide mutational profiles of the HSPCs of the four healthy individuals (HSCT1-4), the patients with mutations in *FANCA* or *FANCC* (PMCFANC01, PMCFANC03, PMCFANC06, PMCFANC08), and the patient with mutations in *BRCA2* (PMCFANC02). **e,** Contribution of base substitution mutational signatures commonly found in blood cells to each FA sample or healthy individual (averaged). Horizontal black lines indicate the expected number of base substitutions based on age. Samples sequenced with bulk WGS are indicated by an asterisk. **f,** Cosine similarities between the mean 96-trinucleotide mutational profiles of the HSPCs of FA patients with the profiles of the healthy HSPCs from the four age-matched donors and the mutational signatures.

**Figure 4.**
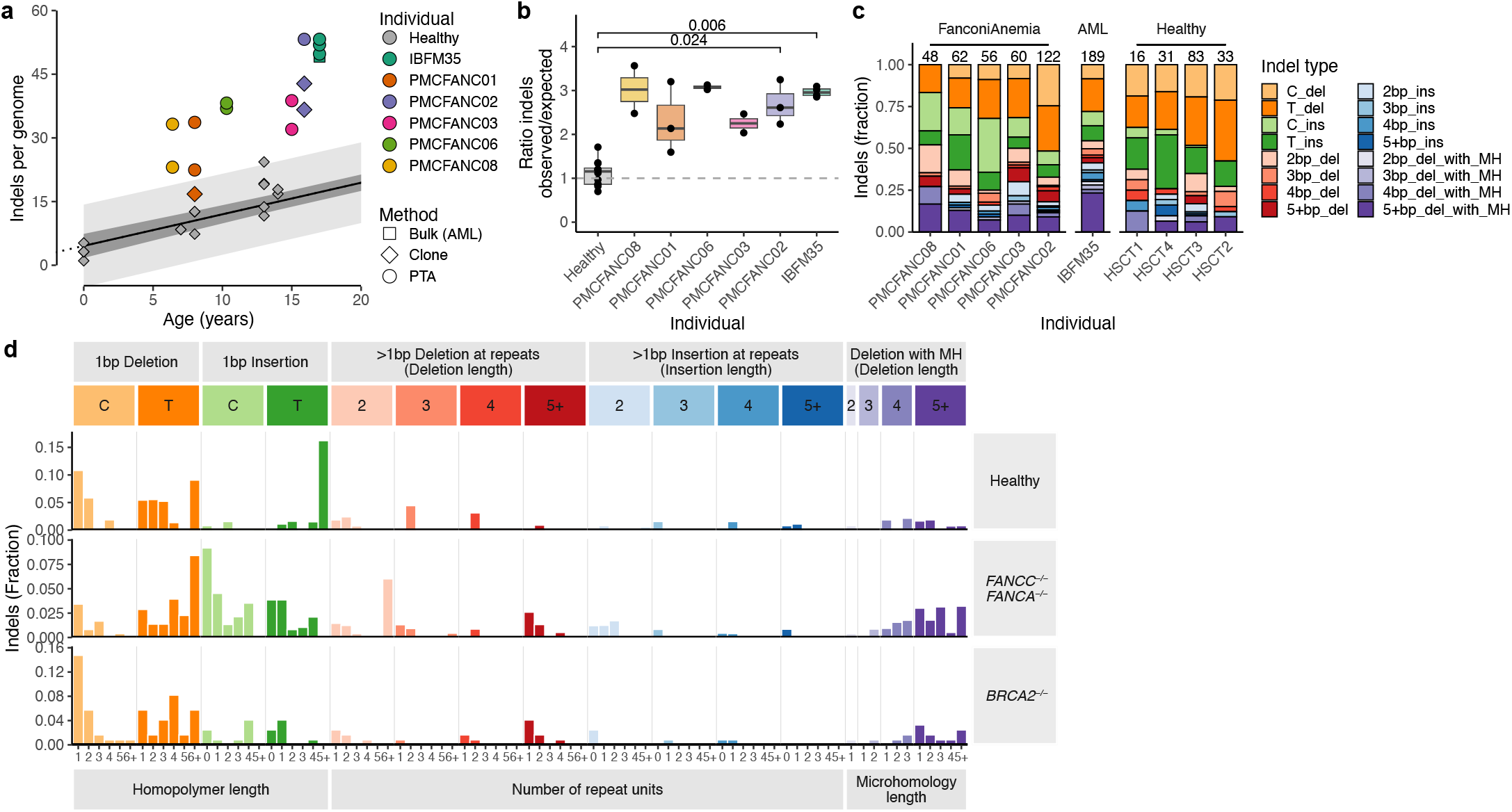
Small insertions and deletions in HSPCs of patients with FA. **a,** Correlation of the number of single base substitutions per HSPC genome of healthy donors (gray points) and patients with FA. Linear mixed modelling showed that healthy HSPCs accumulate indels in a linear fashion with age^21,22^. The 95% confidence interval and the prediction interval of the model are indicated by the dark gray and light gray shading, respectively. **b,** Ratios between the observed and expected number of indels per genome (sorted on age) based on extrapolation of the age linear mixed model. To match the ages of the patients with FA, only 12 HSPCs of four healthy donors (HSCT1-4, ages 7 to 14) are included in this and following panels. P-values indicate multiple testing corrected significant differences (padj<0.05) between three FA patients and the age-matched healthy donors (Bonferroni-corrected Wilcoxon Mann–Whitney test). **c,** Indel spectra showing the relative contribution of the main indel types in the genomes of the donors. Numbers above the bar indicate the total number of indels found in the samples from each individual (without extrapolation). **d,** Total indel profiles of the HSPCs of the four healthy individuals (HSCT1-4), the patients with mutations in *FANCA* or *FANCC* (PMCFANC01, PMCFANC03, PMCFANC06, PMCFANC08), and the patient with mutations in BRCA2 (PMCFANC02).

**Figure 5.**
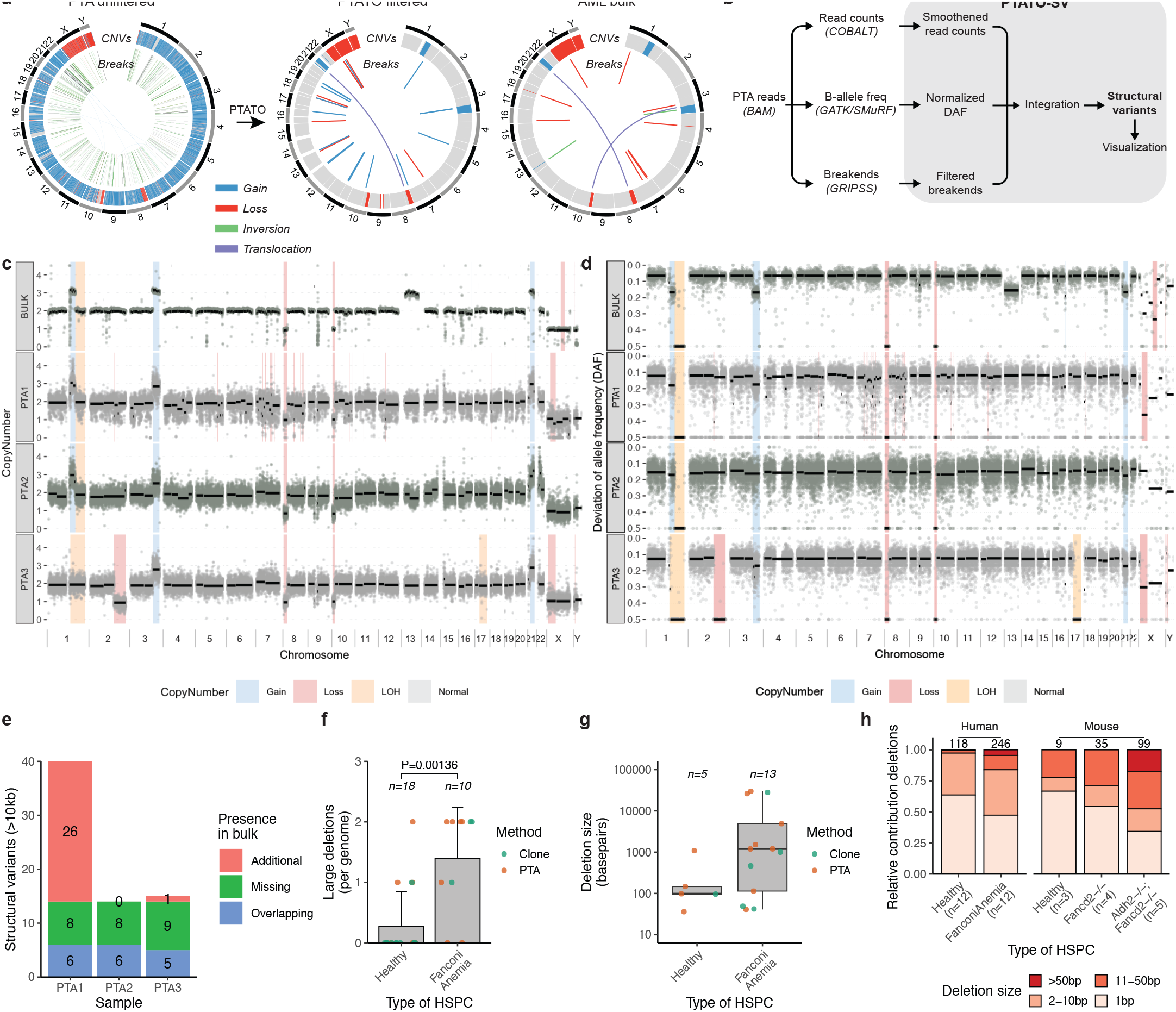
SV filtering by PTATO reveals an increased deletion burden in HSPCs of patients with FA. **a,** Circos plots showing copy number variants (CNVs) and balanced SVs in a PTA (left/center) and bulk WGS sample (right) of patient IBFM35. The standard SV calling pipeline for bulk WGS generates hundreds of false positive calls in PTA samples (left), most of which are removed by PTATO filtering (center), leading to similar SV profiles as a sample sequenced by bulk WGS (right panel). **b,** Schematic overview of the SV calling and filtering strategy tailored for PTA-based WGS data implemented in the PTATO pipeline. **c,** Copy number plots (100kb windows) of the AML-bulk sample analyzed by the bulk-WGS SV calling pipeline and three PTA samples analyzed by PTATO. **d,** Deviation of allele frequency (DAF) plots (100kb windows) of the AML-bulk sample and three PTA samples. The DAF depicts the absolute difference between 0.5 (perfect heterozygosity) and the actual allele frequency of a germline variant. **e,** Number of SVs (>10kb in size) that are present in the HSPCs and present (“Overlapping”) or absent (“Additional”) in the AML-bulk, or present in the bulk but absent in the HSPCs (“Missing”). **f,** Number of deletions (>25bp) detected by GRIDSS and PTATO in genomes of HSPCs of FA patients or healthy donors (including 5 cord blood samples sequenced after PTA). Numbers shown above the bars indicate the number of individuals per group. The P-value was calculated by Wilcoxon Mann–Whitney test. **g,** Size (in bp) of each detected deletion in HSPCs of healthy donors and patients with FA (no significant difference Wilcoxon Mann–Whitney test). Numbers above the boxes indicate the total amount of deletions per group. **h,** Distribution of the sizes of deletions in human and mice^12^ HSPCs with different genetic backgrounds.

The somatic variants detected in the (sub) clones should also be present in the corresponding PTA-amplified samples derived from those (sub) clones and thereby form a reliable set of true positive variants. Between 45-69% of the base substitutions and 31-56% of the indels that were detected in the (sub)clones were also reported in the PTA-amplified cells after PTATO filtering (Figure 2f,g). The clonal variants absent in the PTA-amplified cells were mainly missed due to low coverage and allelic dropout, predominately indicating a limitation of the PTA reaction instead of incorrect filtering by PTATO (Figure 2f,g). Importantly, only 10-16% of the base substitutions and 5-27% of the indels found in both the (sub) clones and the PTA-amplified cells were classified as a PTA artefact by PTATO, showing that PTATO has a mean sensitivity of 86.8% in discriminating true single base substitutions from artefacts (Figure 2f,g). In comparison, SCAN2, a recently developed genotyper for PTA single-cell WGS data^18^, reported on average only 48.8% of the callable variants shared between these PTA-amplified cells and bulk WGS-analyzed (sub)clones (~78% less than PTATO). This finding is in line with the ~46% sensitivity reported for this tool^18^. Indels could not be assessed by SCAN2 for these samples, because it required more PTA samples to build a cross-sample filter list.

We further validated the performance of PTATO by applying it to a previously published PTA-based WGS dataset of human umbilical cord blood cells that were treated with a vehicle (VHC) control or with different dosages of the mutagens D-mannitol (MAN) or N-ethyl-N-nitrosourea (ENU)^17^. Mutational signature analysis showed that filtering by PTATO removed most variants associated with the mutational signature of PTA artefacts, while keeping most single base substitutions associated with signature SBS5 and/or the ENU-associated signature^27^ (Figure S6). These validations show that PTATO can effectively filter single base substitutions and indel artefacts from PTA-based WGS data from different sources, enabling accurate analyses of somatic mutational burdens, patterns, and signatures in single cells.

### Unaltered patterns of indels in most HSPCs of patients with FA

To study the consequences of inactivation of the FA pathway in human HSPCs *in vivo*, we aimed to analyze the genomes of single HSPCs of multiple individuals with FA. However, although we flow sorted at least 200 single HSPCs from six patients for *in vitro* clonal expansion, only for two patients a limited number of clones (one and eight, respectively) expanded to a size large enough for bulk WGS, underlining the need for direct single-cell WGS. Therefore, we used PTA followed by PTATO analysis to study the genomes of single HSPCs derived from bone marrow aspirates of five different individuals with FA (Table 1). In addition, we analyzed the genomes of bulk AML blasts and three PTA-amplified (pre-)leukemic stem cells from a patient with FA (IBFM35) who developed AML after a failed hematopoietic stem cell transplantation.

**Table 1:**
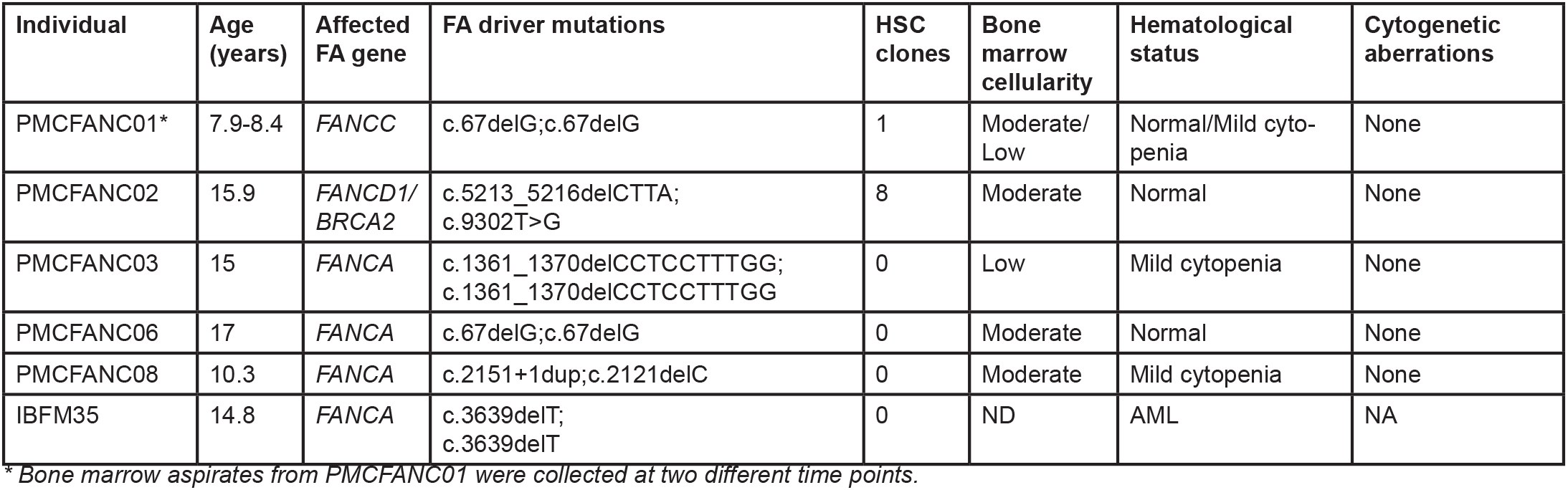
FA patient characteristics at moment of bone marrow puncture.

First, we compared the PTATO-filtered base substitutions detected in the HSPCs of individuals with FA with previously generated WGS data of 34 clonally expanded HSPCs of 11 healthy donors^21,22^. This comparison showed that most of the FA HSPCs had similar somatic single base substitution burdens, patterns and signatures as HSPCs of healthy individuals (Figure 3 and Figure S7). Patient PMCFANC02, whose FA was caused by biallelic germline variants in the *FANCD1/BRCA2* gene, and AML patient IBFM35 formed exceptions with respectively threefold and twofold higher somatic base substitution burden than expected for their age (Figure 3a,b). The elevated mutation burden in PMCFANC02 is mostly caused by base substitutions characterized by signature SBS3, which is associated with homologous recombination deficiency^28,29^, and which is barely detected in the other FA patients (Figure 3c-f).

Subsequently, we compared the somatic indel accumulation between HSPCs of patients with FA and healthy bone marrow donors. The FA HSPCs showed relatively high indel burdens, but only patients PMCFANC02 (*FANCD1/BRCA2*) and IBFM35 had a significantly increased indel burden compared to healthy HSPCs (even in their bulk-sequenced clones and leukemic blasts) (Figure 4a,b). These relatively high indel burdens in FA HSPCs did not seem to be caused by a specific type of indel (Figure 4c,d). These findings, which are in line with observations in FA mouse models^12^ and FA cell lines^24^, confirm that PTATO-based filtering of PTA-based WGS data can be used to accurately study somatic mutations in single cells that cannot be clonally expanded *in vitro*.

### Accurate detection of structural variants in PTA-based sequencing data

It has been shown that HSPCs of FA mouse models and squamous cell carcinomas of human patients with FA have high burdens of somatic structural variants (SVs)^12,30^. Existing bioinformatic tools for single-cell WGS are usually limited to the detection of copy number changes based on read depth^31^ and we found that more comprehensive SV calling pipelines for bulk WGS data detect many false positive variants in PTA-based data (Figure 5a and Figure S8). To study somatic SVs in the HSPCs of the patients with FA, we needed to optimize an SV calling and filtering approach specifically designed for PTA-based WGS data. We used the WGS data of the patient with AML (IBFM35), for who we also have bulk WGS data of AML blasts confirming the presence of different types of SVs, to optimize the SV filtering approach (Figure 5a). PTATO integrates calling of SVs by GRIDSS^32^ and COBALT^33^ based on read depth, B-allele frequencies, split reads and discordant read pairs followed by various normalization and filtering steps tailored for PTA-based WGS data (Figure 5b, Figure S8 and Methods). This approach enables accurate detection of most SVs that were present in the AML bulk sample while reducing the number of false positive calls (Figure 5c). Several SVs present in the leukemic blasts are not detected in the HSPCs, suggesting that these HSPCs are non- or pre-leukemic cells (Figure 5c).

After optimization of SV detection in PTA-based WGS data, we looked for the presence of somatic SVs in the HSPCs of the other patients with FA. We did not observe any large chromosomal abnormalities or translocations (Figure S9). However, we observed 13 deletions with read depth, B-allele frequency (if overlapping germline variants) and split read/discordant read pair support in the 10 cells with sufficient quality ranging from 41 to 29850bp (Figure 5f-h and Supplementary Table S3). The deletions were detected in both the PTA-amplified HSPCs as well as the clonally expanded HSPCs, indicating that the detected deletions are probably not artefacts. Additionally, we rarely observed deletions larger than 100bp in the healthy HSPCs sequenced after clonal expansion or PTA, further supporting that there is an increased burden of deletions in HSPCs of FA patients (Figure 5f-h).

## Discussion

The introduction of PTA greatly improved the accuracy of single-cell WGA, leading to rapid adoption in the field^17,18,34–36^. However, bioinformatic tools making optimal use of the potential of PTA have been lacking. To address this, we developed the PTATO pipeline that can accurately distinguish true positive single base substitutions, indels and SVs from false positive artefacts in PTA-based WGS data. The main benefit of PTATO over other tools, in addition to SV filtering, is the relatively high sensitivity of 86.8% (~78% higher than SCAN2) to distinguish true base substitutions from artefacts. This means that less extrapolation is required to estimate the true somatic mutation burden in cells, which may be especially important for driver mutation detection and retrospective lineage tracing experiments. The RF model included in PTA-TO can be easily retrained if the mutational profiles that are studied are markedly different from the profiles of blood cells that we studied here, making it a flexible tool.

We demonstrated the performance of PTATO by analyzing the genomes of single HSPCs of patients with FA, which could not be clonally expanded *in vitro* for bulk WGS. This analysis showed that most HSPCs of patients with FA have similar somatic mutations burdens as HSPCs of healthy donors, but with an increased number of deletions. These results are in line with findings in mouse models and cell lines of FA^12,24^. The increased deletion burden suggests an increased occurrence of double stranded breaks and/or incorrect repair of these breaks in FA HSPCs, which fits with the molecular functions of the FA DNA repair pathway^8^. It is likely that there is selection against HSCs with more genomic rearrangements without the necessary driver mutations to survive, leading to a gradual depletion of such HSCs in FA patients. The analyzed HSPCs of one FA patient with germline *FANCD2/BRCA2* mutations showed strongly elevated somatic mutation rates, which is consistent with the broader role of BRCA2 independent of the FA DNA repair pathway^37^. This also highlights that the phenotypic heterogeneity between FA patients may be accompanied by genomic heterogeneity in HSPCs between patients^38^. Further studies including larger patient cohorts are required to characterize this genomic heterogeneity, which is likely dependent on the causative germline mutations and disease progression stage.

We showed that our PTATO filtering approach improves the usability of PTA, further narrowing the gap in data quality between single-cell WGS and regular bulk WGS. This will be especially important for the genomic analyses of cells that cannot be clonally expanded for regular WGS, such as diseased or differentiated cells. The accurate characterization of single-cell whole genomes by PTA followed by PTA-TO analysis enables the study of ongoing mutational processes in tissues and cancers, because this combined approach is not limited to analysis of relatively early clonal mutations like regular bulk WGS^39^. We foresee that such single-cell genome analyses made possible by PTATO will yield an unprecedented view of tumor heterogeneity and cancer evolution.

## Methods

### Human bone marrow and umbilical cord blood samples

Bone marrow samples were obtained from the biobank of the Princess Máxima Center for Pediatric Oncology with ethical approval under proposal PM-CLAB2018-007 and PMCLAB2019-027. Written informed consents from the included individuals were obtained by the Princess Máxima Center. The use of material for this study was approved by the Biobank and Data Access Committee of the Princess Máxima Center. The umbilical cord blood sample of donor CB15 was obtained via the University Medical Center Utrecht (UMCU). The collection of cord blood samples was approved by the Biobank Committee of the UMCU (protocol number 19-737). Informed consent for these samples was obtained by the UMCU. The samples from IBFM26 and IBFM35 were obtained from the German Society of Pediatric Oncology and Hematology (GPOH), who also obtained informed consent from these individuals.

### Flow cytometry and primary cell culture

Lin^-^ CD34^+^ HSPCs were single-cell sorted by fluorescence-activated cell sorting (FACS) on an SH800S Cell Sorter (Sony) for clonal expansion or PTA. The following antibodies were used for staining: CD34-BV421 (clone 561, 1:20), lineage (CD3/CD14/CD19/ CD20/CD56)-FITC (clones UCHT1, HCD14, HIB19, 2H7, HCD56, 1:20), CD38-PE (clone HIT2, 1:50), CD90-APC (clone 5E10, 1:200) and CD45RA-PerCP/Cy5.5 (clone HI100, 1:20). AML blasts were selected based on diagnostic immunophenotyping data if available. In most cases, these blasts were CD33, CD38, and/or CD34 positive. All FACS antibodies were obtained from BioLegend.

HSPCs sorted for clonal expansion were cultured in HSPC culture medium for 4 to 7 weeks at 37°C in 5% CO_2_ before collection. HSPC culture medium consisted of StemSpan SFEM medium (STEMCELL Technologies) supplemented with SCF (100 ng/ml), FLT3 ligand (100 ng/ml), IL6 (20 ng/ml), IL3 (10 ng/ml), TPO (50 ng/ml), UM729 (500 nmol/l), and Stemregenin (750 nmol/l). Additionally, mesenchymal stromal cells (MSCs) were cultured from a fraction of bone marrow aspirates by plating cells in 12-well culture dishes with DMEM-F12 medium (Thermo Fisher Scientific) supplemented with 10% fetal bovine serum. The medium was refreshed every other day to remove nonadherent cells, and MSCs could be harvested when confluent (after approximately 2 to 3 weeks).

### Generation of gene knockouts in AHH-1 cell lines

Human B-lymphocyte AHH-1 (CRL-8146) cells were purchased from ATCC. Cells were cultured in RPMI 1640 GlutaMAX medium (Thermo Fisher Scientific) supplemented with 1% Penicillin-Streptomycin (Thermo Fisher Scientific) and 10% horse serum (Thermo Fisher Scientific). Guide RNAs (*FANCC*: 5’-GCAAGAGATGGAGAAGTGTA-3’ and *MSH2*: 5’-GTGCCTTTCAACAACCGGTTG-3’) were cloned into pSpCas9(BB)-2A-GFP (PX458) vector (Addgene#48138). AHH-1 cells were transfected using Lipofectamine 2000 (Thermo Fisher Scientific). One to two days after transfection, GFP-positive transfected cells were single-cell sorted for clonal expansion on a SH800S Cell Sorter (Sony), which was also used for subsequent clonal steps.

MSH2 inactivation was confirmed using western blot, Sanger sequencing and WGS. The following antibodies were used for western blotting: rabbit anti-MSH2 (D24B5, 1:2000, Cell Signaling Technology) and mouse anti-α-Tubulin (T5168, 1:5000, Sigma-Aldrich). Anti-rabbit IgG IRDye 800CW (1:10000, Li-Cor) and anti-mouse IgG IRDye 680RD (1:10000, Li-Cor) were used as secondary antibodies. Western blots were imaged on an Odyssey DLx imaging system (Li-Cor).

FANCC inactivation was validated by Sanger sequencing, WGS and MMC sensitivity assay. For the MMC assay, 5000 cells were plated per well (96-well plates) containing 100μl medium supplemented with different concentrations (0, 5, 10, 50, 100, 500 and 100 nM) of MMC (Sigma-Aldrich) in triplicate. After 5 days of incubation, cell survival was measured using the CellTiter-Glo Luminescent Cell Viability Assay (Promega) according to the manufacturer’s protocol.

For the *MSH2*^-/-^ clonal line, two additional consecutive clonal steps were performed (after 48 and 36 days in culture, respectively), and single cells were sorted for PTA 47 days after the third clonal step (Figure 2a). For the *FANCC*^-/-^ clonal line, a second clonal step was performed 58 days after the first clonal step, and PTA was performed 56 days after the second clonal step (Figure 2a). Four clonal lines were generated for the wildtype cells (Figure 2a). From these four clones, two underwent an additional clonal step (43 and 69 days after the first clonal step) and two were single cell sorted for PTA (84 and 87 days after the clonal step). Cells were harvested for DNA extraction when (sub-)clonal lines were sufficiently expanded after single cell sorts.

### PTA, DNA isolation and WGS

PTA was performed using the ResolveDNA Whole Genome Amplification Kit (BioSkryb Genomics) according to the manufacturer’s protocol. Instead of 10 minutes cell lysis on ice as indicated in the protocol, lysis was performed by 5 minutes incubation on ice followed by 5 minutes incubation at room temperature to maximize DNA denaturation as previously described^34^. DNA samples from bulk AML and bulk MSCs (for germline control) were isolated using the DNeasy DNA Micro Kit (QIAGEN) or DNeasy Blood & Tissue Kit (QIAGEN) according to the manufacturer’s instructions. WGS libraries were generated using standard protocols (Illumina). Libraries were sequenced to 15-30x genome coverage (2×150bp) on an Illumina NovaSeq 6000 system at the Hartwig Medical Foundation (Amsterdam, the Netherlands).

### WGS read alignment and variant calling

WGS reads were mapped against the human reference genome (GRCh38) using the Burrows-Wheeler Aligner (v0.7.17) mapping tool with settings ‘bwa mem –c 100 –M’ ^40^. Sequence reads were marked for duplicates using Sambamba v0.6.8. Realignment was performed using the Genome Analysis Toolkit (GATK) (v4.1.3.0)^41^. A description of the complete data analysis pipeline is available at https://github.com/ToolsVanBox/NF-IAP (v1.3.0). Raw variants were called in multi-sample mode by using the GATK HaplotypeCaller and GATK-Queue with default settings and additional option ‘EMIT_ALL_CONFI-DENT_SITES’. The quality of variant and reference positions was evaluated by using GATK VariantFiltration with options: “--filter-expression ‘QD < 2.0’ --filter-expression ‘MQ < 40.0’ --filter-expression ‘FS > 60.0’ --filter-expression ‘HaplotypeScore > 13.0’ --filter-expression ‘MQRankSum < −12.5’ --filter-expression ‘ReadPosRankSum < −8.0’ --filter-expression ‘MQ0 >= 4 && ((MQ0 / (1.0 * DP)) > 0.1)’ --filter-expression ‘DP < 5’ --filter-expression ‘QUAL < 30’ --filter-expression ‘QUAL >= 30.0 && QUAL < 50.0’ --filter-expression ‘SOR > 4.0’ --filter-name ‘SNP_LowQualityDepth’ --filter-name ‘SNP_MappingQuality’ --filter-name ‘SNP_StrandBias’ --filter-name ‘SNP_HaplotypeScoreHigh’ --filter-name ‘SNP_MQRankSumLow’ --filter-name ‘SNP_Read-PosRankSumLow’ --filter-name ‘SNP_HardToVali-date’ --filter-name ‘SNP_LowCoverage’ --filter-name ‘SNP_VeryLowQual’ --filter-name ‘SNP_LowQual’ --filter-name ‘SNP_SOR’ -cluster 3 -window 10”.

### Processing PTA data from external sources

Single-cell PTA-based WGS data (sra files) from cord blood tissue^17^ were downloaded from the Sequence Read Archive (accession code SRP178894) and extracted into bam files using the prefetch and sam-dump tools of the sratoolkit (v2.9.2)^42^. Samtools view (v1.3) was then used with the “-bf 1” argument to select for the paired reads and Picard SamToFastq (v2.24.1) was used with the “RG_TAG=ID” and “OUTPUT_PER_RG=true” arguments to generate fastq files^40,43^. Seqkit replace (v2.2.0) was used to add a sample id to each read name, because they only consisted of a single read number and a number indicating whether it is the first or second read in the pair^44^. Read alignment and variant calling were then performed as described above.

### PTATO Nextflow implementation

PTATO was implemented in nextflow (v21.10.6.5661). Submodules are containerized and automatically downloaded by a container engine, allowing for an easy installation. Singularity (v3.8.7-1.el7) was used for this manuscript, though Docker will also work with a small change to the config.

### PTATO resources

Next to the sample specific inputs, several general resource files were also used to run PTATO, which are listed in PTATO’s “resources.config” file. To make PTATO easy to install and more reproducible, these resource files are included with downloads of PTATO. First, the fasta file and accompanying indexes of the hg38 version of the human reference genome were downloaded from GATK (https://gatk.broadin-stitute.org/hc/en-us/articles/360035890811). The input files necessary for the COBALT, GRIDSS2, and GRIPSS tools were downloaded from the Hartwig Medical Foundation (https://nextcloud.hartwigmedicalfoundation.nl/s/LTiKTd8XxBqwa-iC?path=%2FHMFTools-Resources)^32,33,45^. A text file containing the centromere locations was downloaded from the UCSC (https://genome.ucsc.edu/cgi-bin/hgTables?hgsid=1424951119_QTS0nx5NshNSyspI7KDoJbVh9tci&clade=mammal&org=Human&db=hg38&hgta_group=map&hgta_track=centromeres&hgta_table=0&hgta_regionType=genome&position=chrX%3A15%2C560%2C138-15%2C602%2C945&hgta_outputType=primaryTable&hgta_outFileName=)^46^. A text file with the genomic coordinates of cytobands was also downloaded from the UCSC (https://genome.ucsc.edu/cgi-bin/hgTables?hgsid=1424951119_QTS0nx5NshNSyspI7KDoJbVh9tci&clade=mammal&org=Human&db=hg38&hgta_group=map&hgta_track=cytoBand&hgta_table=0&hgta_regionType=genome&position=chrX%3A15%2C560%2C138-15%2C602%2C945&hgta_outputType=primaryTable&hgta_outFileName=). A bed file with the genomic coordinates of simple repeats was downloaded from the UCSC for hg19 (http://genome.ucsc.edu/cgi-bin/hgTables?db=hg19&hgta_group=rep&hgta_track=-simpleRepeat&hgta_table=simpleRepeat). A bed file with the genomic coordinates of gene bodies was downloaded from Ensembl for hg19^47^. A bed file with replication timing data was generated as described previously^6^. Files for which hg19 versions were downloaded were converted to hg38 using UCSCs LiftOver tool^46^. Shapeit maps for hg38 were included with Shapeit (v4.2.2)^48^. Shapeit reference haplotype vcf files were downloaded from the 1000 genomes project (http://ftp.1000genomes.ebi.ac.uk/vol1/ftp/data_collections/1000G_2504_high_coverage/working/20201028_3202_phased/).

### Somatic base substitution and indel filtering

The PTATO pipeline uses a multi-sample VCF and sample-specific bam files as input. The somatic variant filtering tool SMuRF (https://github.com/ToolsVan-Box/SMuRF), which is included in the PTATO pipeline, was used to remove germline and low-quality variants by applying several filters as described previously^6^. Briefly, candidate somatic variants were included if they passed the following filters: no evidence in a paired bulk WGS control sample from the same individual; passed by VariantFiltration with a GATK phred-scaled quality score ≥ 100; base coverage of at least 10 (samples with ~30X genome coverage) or 5 (samples with ~15X genome coverage) in the PTA and paired control sample; a mapping quality (MQ) score of >55; and absence of the variant in a panel of unmatched normal human genomes. Additionally, heterozygous and homozygous base substitutions with a GATK genotype score (GQ) lower than 99 or 10, respectively, were removed. Indels with a GQ score lower than 99 in both PTA or paired control sample were removed. Somatic variants with a variant allele frequency of <0.2 were removed.

Variant calling and filtering by SCAN2 was performed using standard settings (including the signature-based rescue step) as described in the manual (https://github.com/parklab/SCAN2/wiki)^18^.

### Allelic imbalance analysis

Before modelling allelic imbalances, variants on each chromosome were phased separately using SHAPEIT (v4.2.2), with the raw vcf file containing all variants as its input^48^. Additionally, the “sequencing” argument was used, SHAPEIT maps for the relevant reference genome were supplied to the map argument and a vcf with reference haplotypes was supplied to the reference argument.

For each candidate somatic variant, first all phased germline variants within 200,000 bp are selected to model allelic imbalance. To ensure only heterozygous germline variants are used, all variants that are not heterozygous in the bulk sample or do not have a dbSNP reference number were removed. After removing all germline variants that were not heterozygous in the sample, the allele depths of all variants phased to the second allele were swapped and the b-allele frequencies were calculated. Next, the b-allele frequencies were fitted with a locally weighted least squares regression, which was used to predict the b-allele frequency of the candidate somatic variant. This regression was performed using the loess R-function with a degree of 2 and using the total allele depth of each variant as weights. Next, a binomial test was performed in R using both the predicted and observed b-allele frequency as well as the total allele depth of the candidate variant, to determine whether the observed allele frequency of the candidate variant matched the surrounding germline variants. The log of the p-value from the allelic imbalance was then used for subsequent steps.

### Selection of sequence context features

For each candidate somatic variant, the surrounding 10bp sequence context and mutation type were retrieved using functions modified from the MutationalPatterns R-package^49^. The “closest” function from bedtools (v2.30.0) was used to identify the genes and simple repeat regions closest to the position of each candidate variant^50^. Bedtools merge (with arguments “-d −1 -o min”) was used to ensure that each mutation is linked to only one feature of each feature list. To identify the transcriptional strand bias and replication timing for each somatic mutation, bedtools was used with the “intersect” argument. Some mutations were linked to multiple overlapping gene annotations. For the transcriptional stand bias this was solved by using bedtools with the “merge -d −1 -o distinct” arguments to check if a variant was present in the plus strand, minus strand or both. For the replication timing bedtools was used with the “merge -d −1 -o median” arguments to merge mutations that are present in multiple genes. Next, to merge the gene body, simple repeat, transcriptional strand bias, and replication timing features for each variant, bedtools was used with the “intersect” argument, after which the variants were merged using bedtools with the “merge -d −1 -o unique” arguments.

### Linked read analysis using read-backed phasing

For each heterozygous candidate somatic variant, all sequencing reads overlapping the position of the variant were extracted from the sample’s bam file. Additionally, all heterozygous germline variants within the area spanned by the reads are extracted from the original input vcf. Next, for each germline variant each read that spans both the germline and somatic variant is checked. Each read that contains either the alternative alleles for both the germline and somatic variant or the reference alleles for both the germline and somatic variant is counted as a cis read. Other reads are counted as trans reads. If a candidate is real, then it would be expected that almost all reads are either cis or trans. Whether the variants are cis, trans, or mixed is then calculated based on a Bayesian likelihood score similar to the one used by SV-Typer^51^. The likelihood scores of the three options are then combined into a single Phred-scaled quality score. Candidate variants with a score of <100, between 100-1000 and >1000 were considered to be false positive, uncertain or true variants, respectively.

### Random forest training

To obtain a set of true positive variants for training the RF model, base substitutions were selected that were detected in PTA samples of IBFM26, IBFM35, PB10268 and PMCAHH1-FANCCKO and also in bulk WGS-analyzed samples from the same individuals (Figure 1b and Supplementary Table S1). Somatic base substitutions with a linked read analysis score below 1 in these samples were included in the set of artefacts. Variants that were shared between PTA and bulk WGS samples and also had a linked read analysis score of less than 1 were excluded from both the true positive and the artifact datasets. Variants overlapping with copy number variants and regions of loss-of-heterozygosity in samples of IBFM26, IBFM35 and PMCAHH1-FANCCKO were excluded from training. Additionally, unique base substitutions detected in three umbilical cord blood HSPCs of donor PMCCB15 analyzed by PTA were considered artifacts, as the number of true mutations in the cord bloods is expected to be very low (20-50)^21^. Finally, the number of base substitutions in the artefact set was subsampled to be the same as the number of base substitutions in true positive set to result in a better class balance.

A random forest was trained on the previously described features with the randomForest (v 4.7-1) R package supplying the “mtry” argument with a value of 4. For some variants no p-value for the allelic imbalance or no replication timing value could be calculated, therefore they were excluded from the training. Instead, two more random forests were trained that did include these variables. One without the allelic imbalance variable and one without both this variable and the replication timing variable.

### Candidate variant classification by PTATO

For each candidate somatic base substitution, PTATO’s RF model calculated a probability score to predict if a variant is a PTA artefact. A higher score indicates a higher probability that a variant is an artefact according to the RF. Subsequently, two methods were used to determine a sample-specific cutoff value (variants above the cutoff were considered to be artefacts).

First, for each sample a group of likely true positive variants and a group of likely artefacts were selected by taking the variants with either a high (>=1000) or low (<1) linked read analysis score. These variants classified by the linked read analysis were used to validate the performance of the RF model. Precision and recall were calculated for a range of prediction score cutoff values (between 0 and 1 with increments of 0.01). The optimal linked read analysis cutoff was determined by taking the insersection of the precision-recall curves.

Second, a range of different cutoff values (from 0.1 to 0.8 with increments of 0.025) was taken and for each of these cutoffs the variants with a probability score below the cutoff were selected (leading to 29 groups of mutations). For all these 29 groups of mutations, a 96-trinucleotide mutation matrix was calculated using MutationalPatterns. Subsequently, the cosine similarities between all those groups were calculated using the calc_cosim_mutmat() function from MutationalPatterns. Hierarchical clustering of the cosine similarities was performed using the hclust() function in R (Euclidean distance with complete linkage) to generate two clusters: one cluster with low PTA probability cutoffs (and mostly true positives) and one cluster with relatively high cutoffs (and mostly false positives). The highest cutoff value in the cluster with true positives was taken as the cosine similarity cutoff.

Finally, the linked read analyses cutoff and cosine similarity cutoff were merged into a final cutoff that was used to classify variants as true or false positive. This was done by taking the mean of both cutoffs, or by only selecting the cosine similarity cutoff if the highest precision-recall value of the linked read analysis cutoff was below 0.7 (for example in case there were too few variants classified by the linked read analysis).

### Indel filtering

Candidate indels were filtered based on recurrency in multiple unrelated individuals. Raw indel calls from 31 PTA-based WGS samples from four unrelated individuals were merged into one vcf using bcftools (v1.9). Indels detected in samples from at least two different individuals were included in the PTATO indel exclusion vcf file. Candidate indels present in the exclusion vcf file are removed from test samples. Additionally, insertions in 5bp+ homopolymers were removed. For this, MutationalPatterns was used to determine the indel type and sequence context around candidate indels.

### Mutation burden and signature analysis

The mutational patterns and signature analyses were performed using MutationalPatterns (v3.6.0)^49^. Mutational signatures were used from COSMIC (v3.2) as well as the previously described HSPC, PTA, and ENU signatures^18,21,27,52^. Figures were made using ggplot2 (v3.4.0)^53^.

CallableLoci from GATK v3.8.1 (with parameters --minBaseQuality 10 --minMappingQuality 10 --minDepth 8 --minDepthForLowMAPQ 10 --maxDepth 100) was used to determine the fraction of the sequenced genome that had sufficient coverage and quality for variant calling. Variants not overlapping with the callable regions determined by CallableLoci were excluded. Subsequently, all remaining variants on autosomal chromosomes were counted. To obtain the mutation burden, the mutation count was extrapolated by dividing it by the fraction of the genome that was surveyed (determined by CallableLoci), as previously described^6^.

A linear mixed-effects model was used to correlate the mutation burden in HSPCs from healthy donors and the age of the donors as previously described^22^. This model was used to calculate the expected mutation burdens for the specific ages of the patients. The 95% confidence and 95% prediction intervals were calculated using the R package ggeffects (v1.1.0)^54^.

### Normalization of copy number ratios for SV detection

GC-normalized read depth per 1000 basepair genomic window was calculated by COBALT (v1.11)^33^. A coverage panel-of-normals (PON) was generated by merging COBALT ratio files of 12 copy number neutral PTA samples. The total read counts from all windows of each sample were first normalized so that every sample has the same total amount of read counts. Subsequently, the mean readcount per bin over all normal samples in the PON was calculated. PTATO uses the coverage PON file to smoothen PTA-specific coverage fluctuations. First, the total read depth in a test sample is normalized to the same total amount of read counts in the coverage PON. Subsequently, the read counts in each window are divided by the mean read counts in the same window in the PON. Additionally, the bottom and top 1% outlier windows in the PON file and the windows located within 1Mb distance of centromeres and telomers are excluded from the analysis.

The smoothened read counts were subsequently binned in 100kb windows. The copynumber (v1.34.0) R-package with parameter “gamma=100” was used to segment the median read count data in both the 100kb and 1kb windows^55^. The segments based on the 100kb resolution were used as raw copy number segments. The start and end coordinates of these raw copy number segments were fine mapped by taking the start and end coordinates of overlapping 1kb window-based segments. Fine mapped segments with a copy number ratio of <1.5 were considered to be copy number losses and segments with ratios >2.5 were considered to be copy number gains.

### Deviation of allele frequency calculations

Variant allele frequencies (VAF) of germline variants can be noisy in PTA-based WGS data due to uneven genome amplification, which impedes accurate copy number variant detection based on raw B-allele frequencies. To reduce noise due to uneven amplification, the VAFs of germline base substitutions were first binned in 100kb windows instead of taking separate B-allele frequencies of each individual variant. To determine a mean allele frequency for multiple variants in a bin, the deviation of allele frequency (DAF) was calculated by taking the absolute value after subtracting the VAF of each variant from 0.5 (which is the expected VAF for a perfectly amplified and sequenced germline variant). Thus, each variant has a DAF between 0 (corresponding to a VAF of 0.5) and 0.5 (corresponding to a VAF of 0 or 1). Subsequently, all DAF values of germline base substitutions are binned in 100kb genomic regions and the mean DAF for each region is calculated. The copynumber R-package with parameter “gamma=100” was used to segment the 100kb bins in crude DAF regions. These crude segments were fine mapped by adjusting the start and end coordinates of the segments to the positions of the nearest germline SNVs (within 200kb distance of the segment) with similar DAFs as the segment. Segments with a DAF of more than 0.4 (corresponding to VAF < 0.1 or > 0.9) were considered to be loss of heterozygosity regions. Segments with a DAF between 0.16 and 0.4 (corresponding to VAFs between 0.1 to 0.32 or 0.64 to 0.9) were considered to be regions with copy number gains.

### SV breakend calling and filtering

Somatic SV breakends were called by GRIDSS v2.13.2 and prefiltered by GRIPSS v1.9 using a corresponding bulk-sequenced germline control^32^. The GRIPSS-filtered somatic breakends of 15 PTA-based samples of four unrelated individuals were merged using bedtools merge (v2.30.0). Breakend positions occurring within 2000bp of each other in multiple of these individuals were included in a breakend PON. Candidate breakends in other samples overlapping with the regions in the breakend PON were removed. Subsequently the normalized coverage and DAF of the SV candidates was calculated. Breakends of duplications were filtered if the DAF was less than 0.18 and/or the copy number ratio was <2.5. Breakends of deletions were filtered if the DAF was less than 0.4 and/or the copy number ratio was >1.5. Breakends with a coverage of more than 100 were also excluded for samples with a targeted genome coverage of 15x as many artefacts occur in these regions with excess coverage. Inversions were filtered if they only have one breakpoint junction instead of two. Additionally, all inversions less than 1kb in size were removed. Inter-chromosomal events were also filtered if they only have one breakpoint junction (instead of two), unless they were situated less than 100kb from a copy number variant. This exception rescues unbalanced translocations.

### Integration of coverage, allele frequencies and structural variant breakends

The coverage segments, DAF segments, and breakends of SV candidates were intersected to create the final list of filtered structural variants. Copy number changes were required to have both coverage and DAF support, but not necessarily breakend support, as many CNVs have start and/or end positions within repeat regions that are difficult to capture with PTA and/or short-read sequencing. Regions with a DAF >0.4 (corresponding to VAFs of <0.1 and >0.9) without coverage support (copy number >1.5) were considered to be loss-of-heterozygosity regions. ggplot2 and Circos (v0.69-9) were used for to visualize structural variants and karyograms^56^.

## Data and code availability

Source code and manual for PTATO are available at https://github.com/ToolsVanBox/PTATO/

## Acknowledgments

We are grateful to the donors for their participation in this study. We would like to thank Agnes Vissers, Edwin Sonneveld and the biobank of the Princess Máxima Center for their assistance with inclusion of the donors. We thank the Hartwig Medical Foundation (Amsterdam, the Netherlands) for facilitating WGS. This research was supported by a grant from the Dutch Cancer Society (KWF Research Project 12682) to R.v.B and The New York Stem Cell Foundation. R.v.B. is a New York Stem Cell Foundation – Robertson Investigator.

## Author contributions

S.M., M.E.B. and R.v.B. conceived and designed the study. S.M., F.M. and M.v.R. developed the PTATO computational pipeline. S.M. and F.M. performed computational analyses and designed the figures. F.P., E.J.M.B., I.v.d.W., L.L.M.D., L.T., A.M.B. and S.M. performed sample collection and single cell isolations using flow sorting. D.M.G. performed SCAN2 variant filtering. N.M.G, M.V., L.T. and S.M. generated the AHH-1 cell lines used in this study. S.M., L.T. and M.V. performed PTA. M.E.B., M.B., E.A., D.R. arranged inclusion of the donors and collection of donor material. The manuscript was written by S.M, F.M. and R.v.B. with contributions from all authors.

