## Supplemental Figures for "Comprehensive single-cell genome analysis at nucleotide resolution using the PTA Analysis Toolbox"

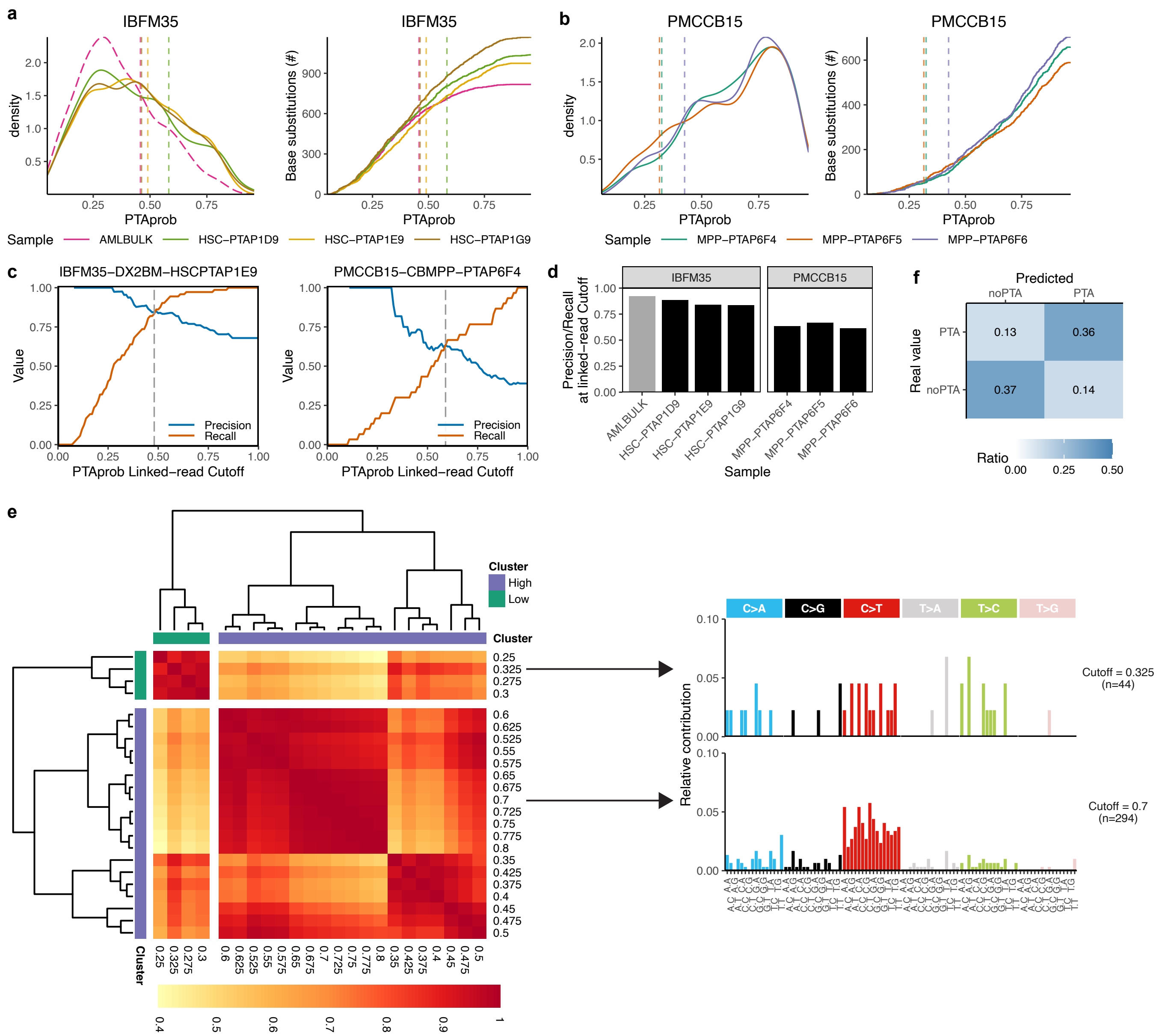

**Figure S1. Calculations of optimal PTA probability cutoffs by PTATO. a,** Distributions (left) and cumulative distributions (right) of the PTA probability scores (PTAprob) of candidate base substitutions before PTATO filtering in one bulk WGS (with relatively low PTAprob scores) and three PTA-based WGS samples. Vertical lines indicate the sample-specific PTAprob cutoffs determined by PTATO. **b,** Same as a, but then for umbilical cord blood samples with low mutation burdens. **c**, Precision and recall at different PTAprob cutoffs of a subset of base substitutions that could be classified as true or false positive by the linked read analysis. The linked read cutoff is determined by taking the PTA probability at minimal difference between the precision and recall. **d**, Overview of the linked read precision-recall rates of samples in the training set. Samples with low mutations burdens can have low precision-recall rates, as shown here for cord blood donor PMCCB15, which requires an alternative method to calculate an optimal cutoff. **e,** Heatmap showing the cosine similarities between 96-trinucleotide mutational profiles calculated for different PTAprob cutoffs in sample PMCCB15-CBMPP-PTAP6F4. Hierarchical clustering is used to make one cluster with low PTAprob cutoffs (containing most true positives) and one cluster with high PTAprob cutoffs (containing most artefacts). The highest value in the cluster with true positives is used as the cosine similarity cutoff (0.325 in this case). Two example profiles of the mutation sets at different cutoffs are shown on the right. **f,** Heatmap depicting the ratio of true positives, false positives, true negatives and false negatives for single base substitutions in the training set classified by PTATO.

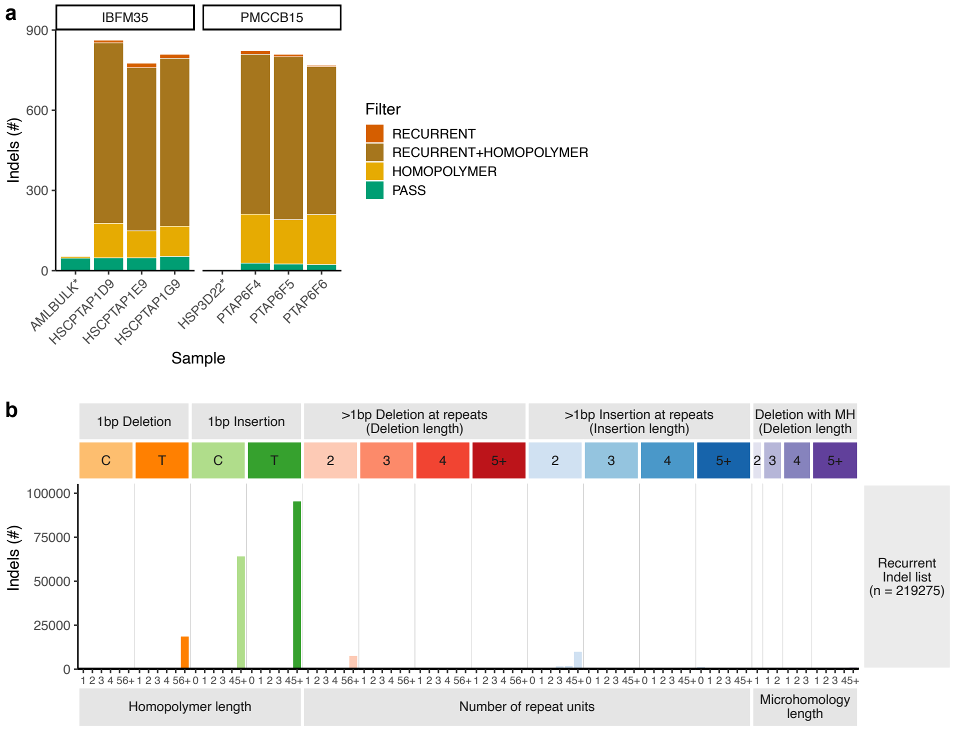


**Figure S2. Indel filtering by PTATO based on recurrency and sequence context. a,** PTATO filters indel artefacts by filtering insertions at long homopolymers (HOMOPOLYMER) and by filtering indels recurrent in multiple unrelated individuals (RECURRENT). This filtering removes most excess indels (the remaining indels are labelled with PASS), but also limits sensitivity to detect insertions in long homopolymer tracts. **b,** Profile of the indels present in the list of recurrent indels that is used by PTATO to filter indel artefacts. The exclusion list contains mostly insertions at long homopolymers, but also some recurrent deletions at long homopolymers. This indicates that just excluding insertions at long homopolymers is not sufficient to remove all indel artefacts.

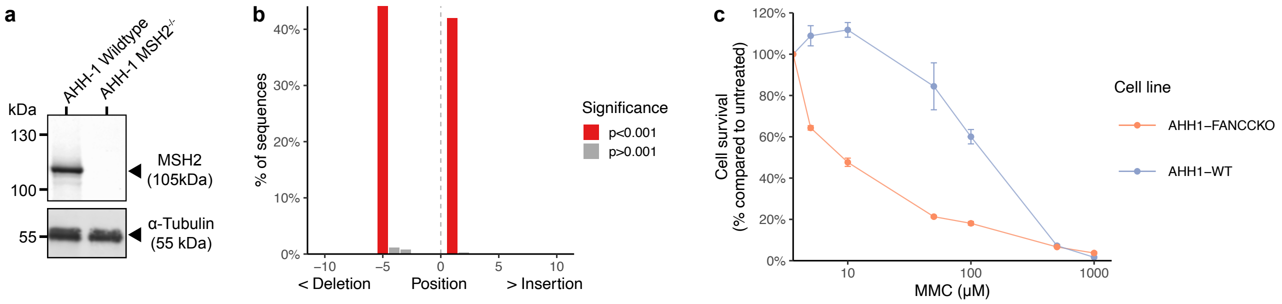


**Figure S3. Validation of *MSH2* and *FANCC* knockout status in AHH-1 cell lines. a,** Western blot showing the absence of MSH2 protein expression in the AHH-1 *MSH2*$^{-/-}$ clonal cell line. **b,** TIDE analysis detects a 5-basepair deletion and 1-basepair insertion introduced by CRISPR/Cas9 in the *FANCC* gene of the AHH-1 *FANCC*$^{-/-}$ clonal cell line. Due to the absence of high quality antibodies, western blotting could not be performed to study FANCC protein expression. Therefore, we used PCR and Sanger sequencing followed by TIDE decomposition, in addition to a Mitomycin C (MMC) sensitivity assay, to confirm knockout status. The presence of the biallelic indels in *FANCC* was also confirmed in the WGS data (data not shown). **c,** MMC sensitivity assay showing the hypersensitivity of the AHH1 *FANCC*$^{-/-}$ clonal cell line to the DNA cross-linking agent MMC. This finding provides additional support for the knockout status of *FANCC* in this cell line, as cells of patients with FA are known to display MMC hypersensitivity. Mean survival values from triplicate experiments are shown and error bars indicate standard deviations.

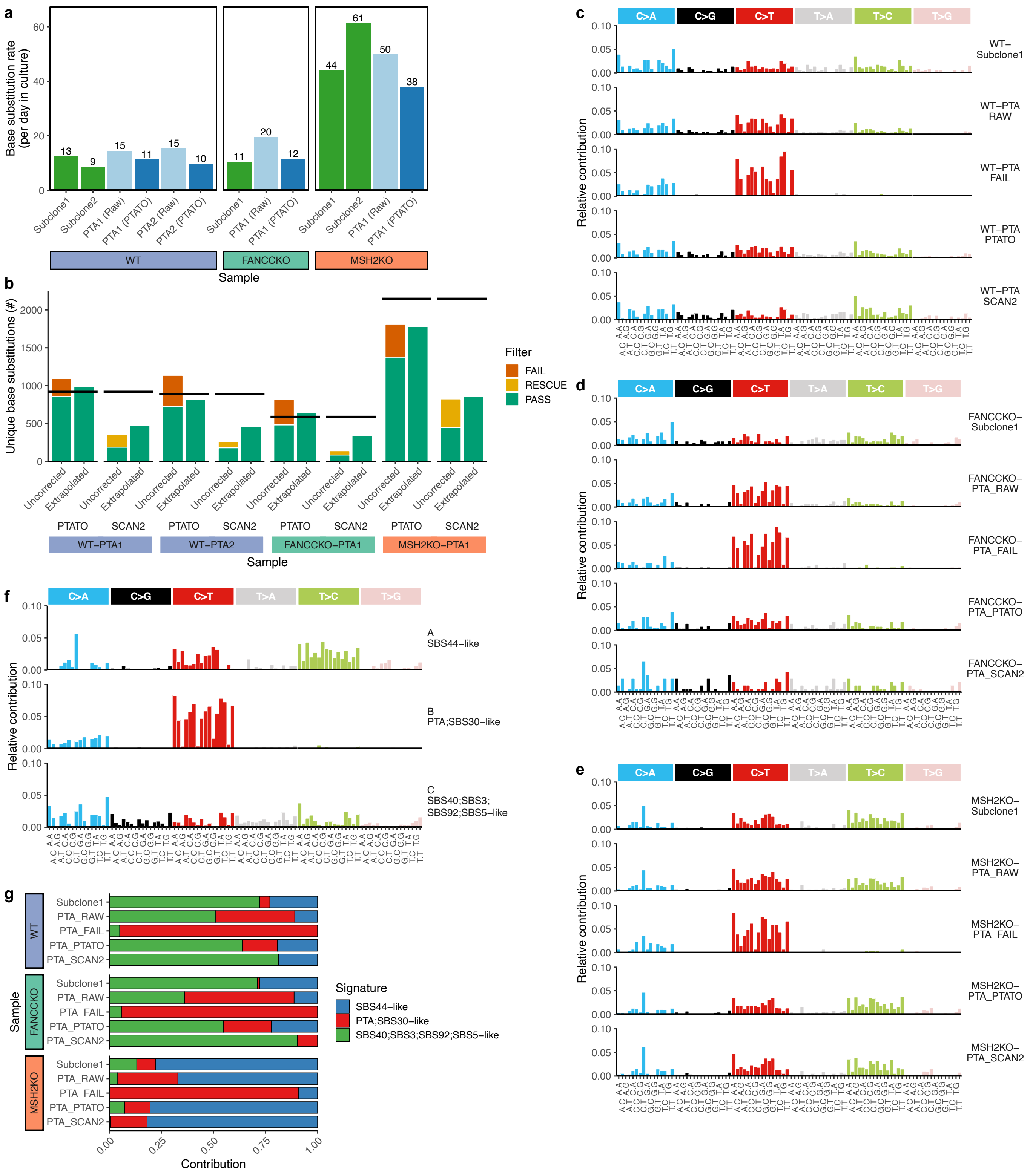

**Figure S4. Single base substitution filtering in PTA-based WGS data of AHH-1 cell lines by PTATO. a**, Number of base substitutions aqcuired per day in culture between single cell steps in subclones analyzed by bulk WGS (green) and PTA samples before (lightblue) and after (darkblue) PTATO filtering. **b,** Number of unique base substitutions not present in the (sub)clones reported by PTATO and SCAN2 before and after extrapolation. PTATO detects more base substitutions, requiring less extrapolation to estimate the true base substitutions burden in a cell. The horizontal black lines indicate the expected number of base substitutions based on the days in culture since the previous single-cell step and the mutation rate in the corresponding subclones. **c-e,** The 96-trinucleotide mutational profiles of the wildtype (WT) (**c**), FANCC-KO (**d**) and MSH2-KO (**e**) AHH-1 cells assesed by WGS after clonal expansion or after PTA. The variant calls before PTATO filtering (RAW) still contain numerous PTA artefacts. The profiles of the variants removed by PTATO are shown in the middle panels (PTA_FAIL). **f**, 96-trinucleotide profiles of the base substitution signatures extracted by non-negative matrix factorization (NMF). One signature resembles the PTA artefact signature (red), one resembles the background signature for AHH-1 cells (green) and one resembles signatures found in mismatch repair deficient cells (blue). **g,** Contribution of the signatures extracted by NMF (f) to the mutational profiles of each sample. The mutations removed by PTATO (PTA_FAIL) are mostly refitted to the PTA artefact signature. The mutational profiles of the PTA samples filtered by PTATO and SCAN2 are more similar to the profiles of the subclones analyzed by bulk WGS than the unfiltered (PTA_RAW) samples.

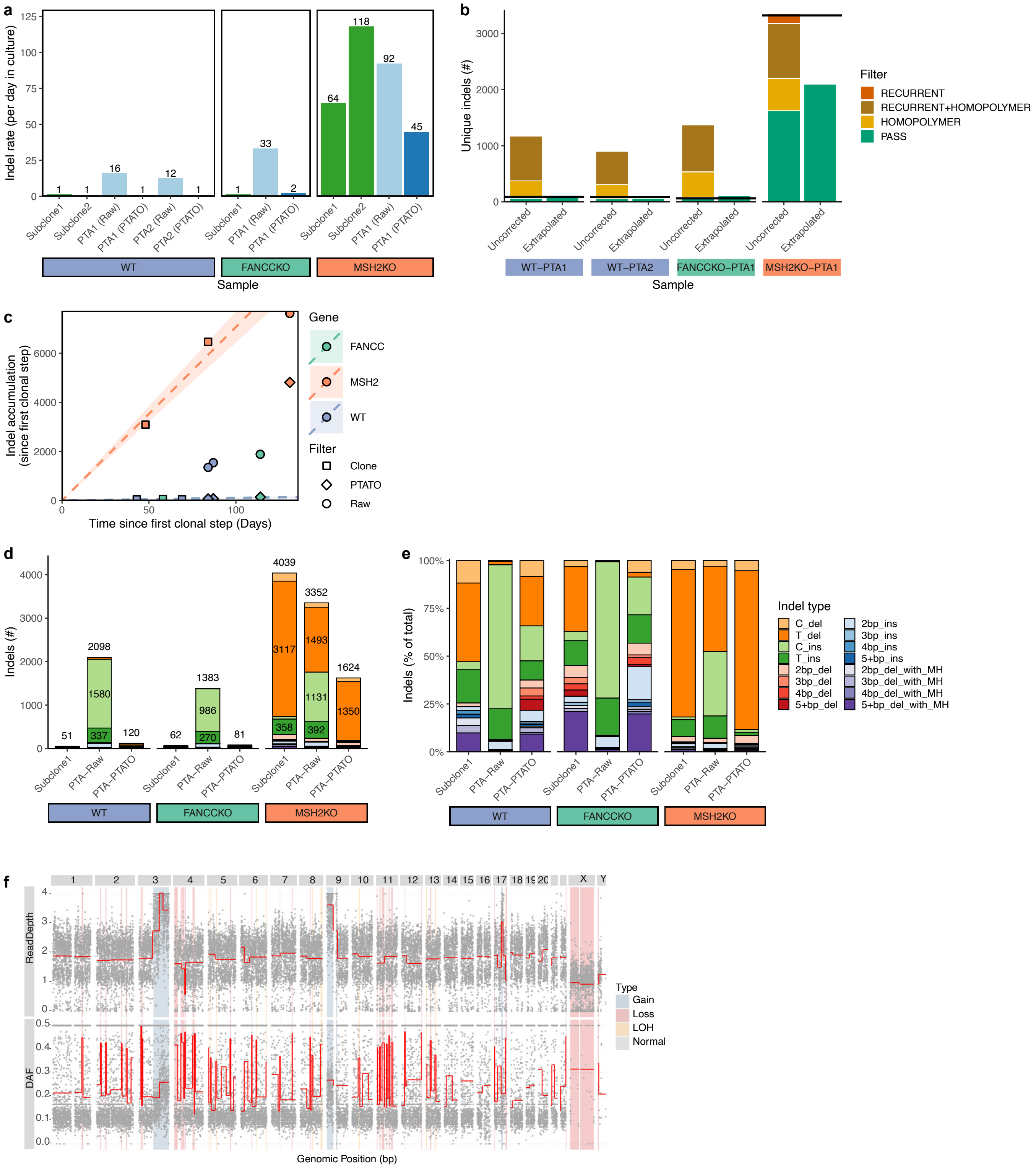


**Figure S5. Indel filtering in PTA-based WGS data of AHH-1 cell lines by PTATO. a**, Number of indels aqcuired per day in culture between single cell steps in subclones analyzed by bulk WGS (green) and PTA samples before (lightblue) and after (darkblue) PTATO filtering. **b,** Number of unique indels not present in the (sub)clones reported by PTATO before and after extrapolation. The horizontal black lines indicate the expected number of indels based on the days in culture since the previous single-cell step and the indel accumulation rate in the corresponding subclones. **c,** Accumulation of indels since the first clonal step. The circles and diamonds indicate the number of indels detected in the PTA samples before and after PTATO filtering, respectively. **d,** Number of indels (not present in the preceding clonal step) in the subclones analyzed by bulk WGS and the PTA samples before (Raw) and after PTATO filtering. More than a thousand arteficial indels are detected in the wildtype and FANCC$^{-/-}$ PTA samples. **e,** Relative contributions of the different types of indels (not present in the preceding clonal step) detected in the subclones analyzed by bulk WGS and the PTA samples before (Raw) and after PTATO filtering. PTATO mostly removes 1-basepair (bp) insertions. **f,** Copy number and deviation-of-allele frequency plots of sample PMCAHH1-MSH2KO-C27E06SC51B06-PTAP1E7. This sample has many loss-of-heterozygosity (LOH) regions, indicating a lower quality genome amplification by PTA. MH, microhomology.

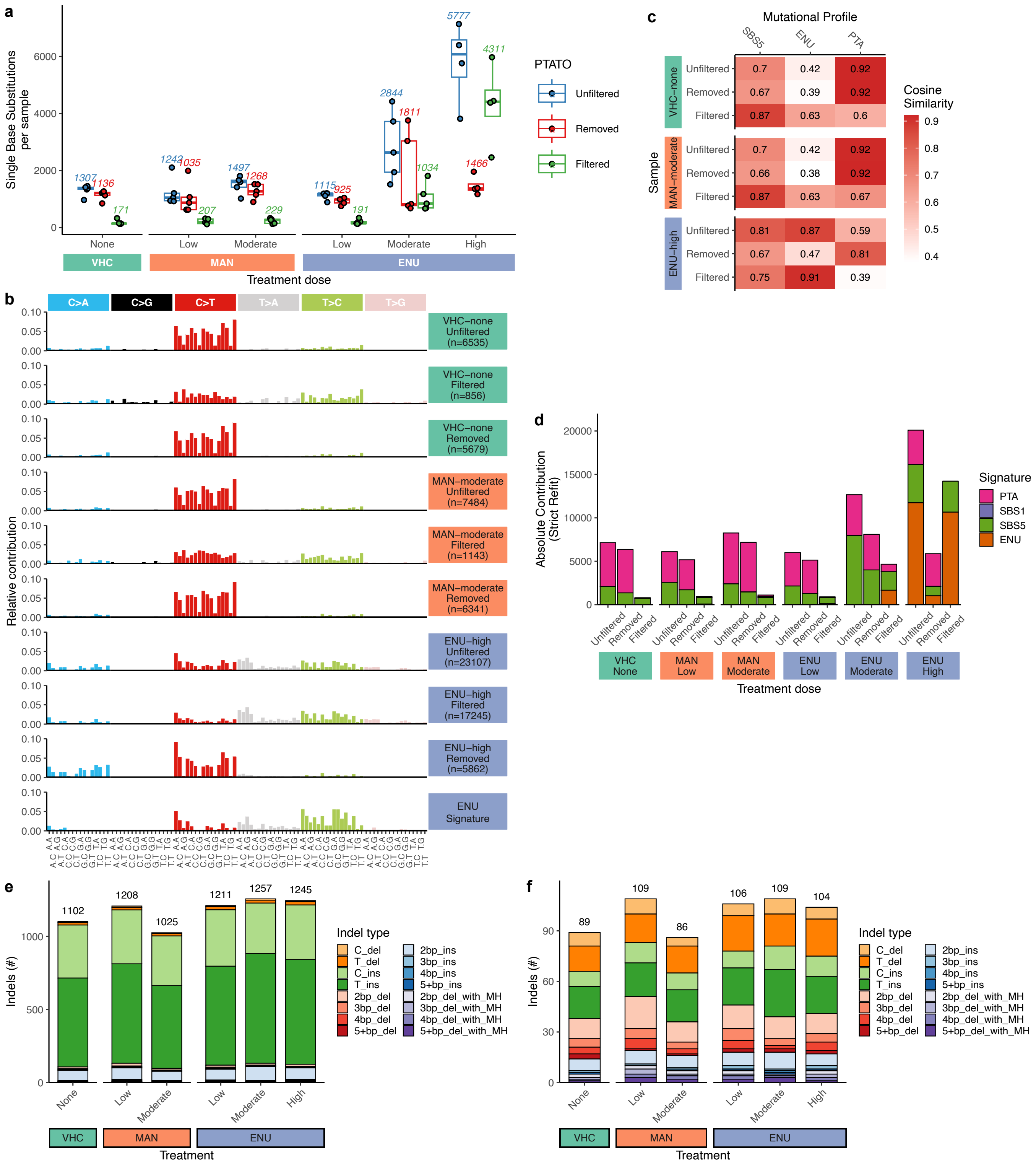


| Sample | | SBS5 | ENU | PTA |
|---|---|---|---|---|
| VHC–none | Unfiltered | 0.7 | 0.42 | 0.92 |
| | Removed | 0.67 | 0.39 | 0.92 |
| | Filtered | 0.87 | 0.63 | 0.6 |
| MAN–moderate | Unfiltered | 0.7 | 0.42 | 0.92 |
| | Removed | 0.66 | 0.38 | 0.92 |
| | Filtered | 0.87 | 0.63 | 0.67 |
| ENU–high | Unfiltered | 0.81 | 0.87 | 0.59 |
| | Removed | 0.67 | 0.47 | 0.81 |
| | Filtered | 0.75 | 0.91 | 0.39 |

**Figure S6. PTATO accurately filters PTA artefacts from a PTA-based WGS dataset of umbilical cord blood cells. a,** Boxplot showing the number of base substitutions for human cord blood samples treated with different concentrations of a vehicle control (VHC; n = 5), D-mannitol (MAN; low: n = 5, moderate: n = 5) or N-ethyl-N-nitrosourea (ENU; low: n = 5, moderate: n = 5, high: n = 4). Numbers indicate the mean base substitution burden per sample in each treatment group. **b,** The 96-trinucleotide profiles of the base substitutions in the indicated treatment groups before ("Unfiltered") or after ("Filtered") PTATO filtering or the base substitutions removed by PTATO ("Removed"). The bottom panel shows the profile of the mutational signature that has been previously associated with ENU-treatment. **c,** Cosine similarities of the mutational profiles of the base substitutions that are present before ("Unfiltered") or after PTATO filtering ("Filtered"), or that are removed by PTATO, with the SBS5-, ENU- and PTA mutational signatures. **d,** Contribution of the PTA, SBS1, SBS5 and ENU mutational signatures to the profiles of the unfiltered, removed and filtered base substitutions determined by a strict mutational refit. PTATO mostly removes mutations associated with the PTA mutational signature, while keeping the mutations associated with SBS5 and the ENU mutational signatures. The base substitutions were pooled for each treatment dose. **e,** Mean numbers and types of indels found per sample in each treatment group before filtering by PTATO. **f,** Mean numbers and types of indels found per sample in each treatment group after filtering by PTATO. PTATO removes over a thousand indels per sample, mainly C- and T-insertions at homopolymers. As has been shown before, treatment with ENU did not cause an increase in indel burden. MH, Microhomology.

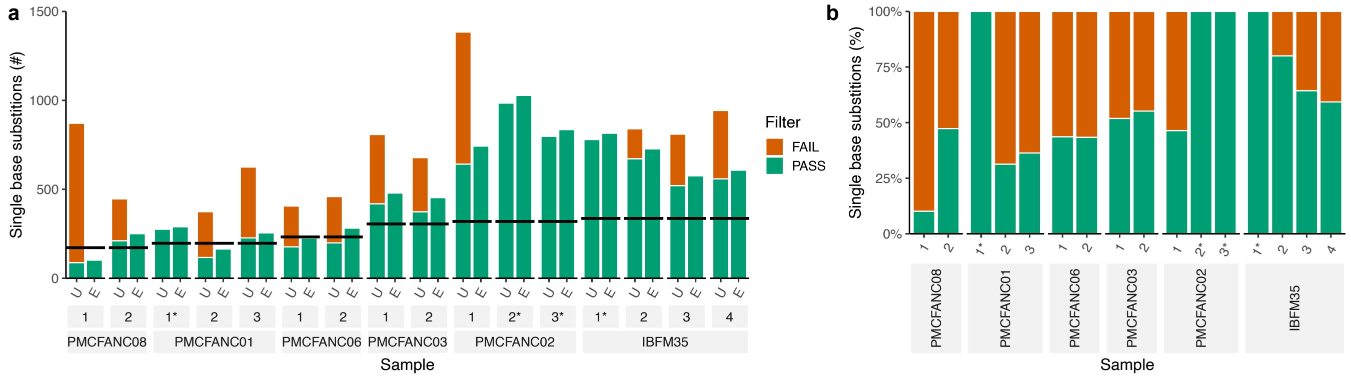


**Figure S7. PTATO filtering of single base substitutions in HSPCs of patients with FA. a,** Absolute number of single base substitutions that passed (PASS) or failed (FAIL) filtering by PTATO, before (U = unfiltered) and after (E = extrapolated) extrapolation based on CallableLoci. The horizontal lines indicate the expected number of base substitutions for each individual based on their age. Samples not amplified by PTA are marked with an asterisk. **b,** Relative amount of single base substitutions that that passed or failed filtering by PTATO.

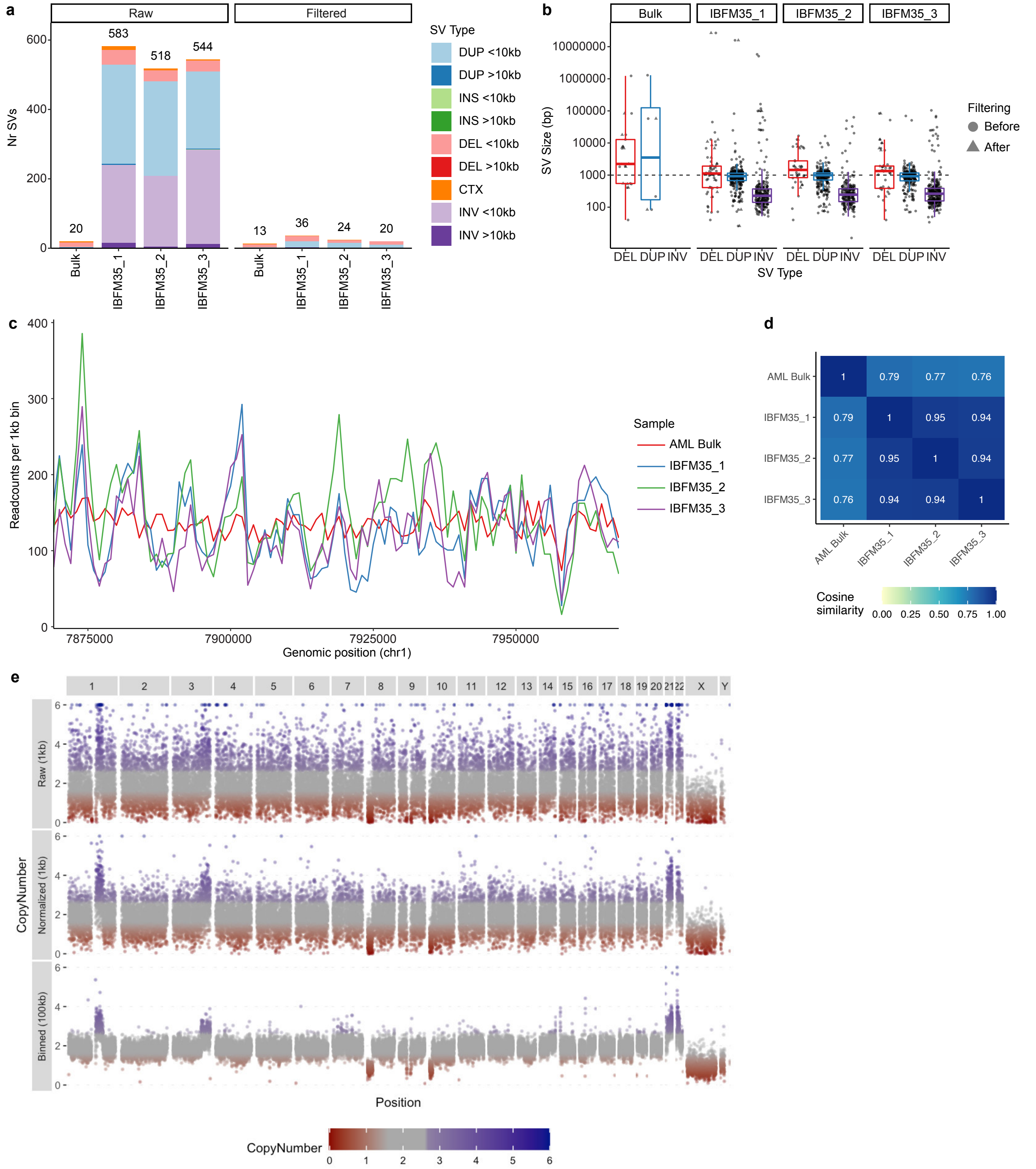


**Figure S8. SV filtering by PTATO. a,** Number and types of SVs detected before ("Raw") and after PTATO filtering in AML blasts (bulk-WGS) and three single HSPCs (PTA-based WGS) of AML patient IBFM35. GRIDSS calls many false positive SVs in PTA-based WGS data, predominantly inversions and duplications <10 kilobases (kb) in size (left). Most of these false positive SVs are removed by PTATO (right). **b,** Sizes (in bp) of the deletions (DEL), duplications (DUP), and inversions (INV) in the bulk sequenced AML sample and three PTA-based HSPC samples from patient IBFM35 before and after PTATO filtering. **c,** Normalized sequencing read counts per 1kb bin (determined by COBALT) in a 100-kb region of chromosome 1 in the bulk sequenced AML sample and three PTA-based HSPC samples from patient IBFM35. The coverage in the PTA samples follow similar patterns that are different and noisier compared to the bulk-WGS sample. **d,** Heatmap depicting the cosine similarities of read counts in 1kb bins between the AML sample analyzed by bulk WGS and three HSPCs analyzed by PTA-based WGS. It shows that the coverage patterns are very similar between PTA-based WGS samples. These recurrent patterns make it useful to smoothen the coverage in PTA samples by normalizing it against other PTA samples. **e.** Example of copy number smoothening by PTATO for sample IBFM35_2. The top panel shows the GC-normalized copy number per 1kb bin determined by COBALT, which is relatively noisy for PTA-based WGS data. Subsequently, the copy numbers per 1kb bin are normalized against a panel of other PTA samples, reducing part of the noise as shown in the middle panel. Finally, the normalized read counts are binned in 100kb windows as shown in the bottom panel. These read counts in 100kb windows are used for segmentation and copy number variant calling. INS = Insertion; CTX = Inter-chromosomal translocation

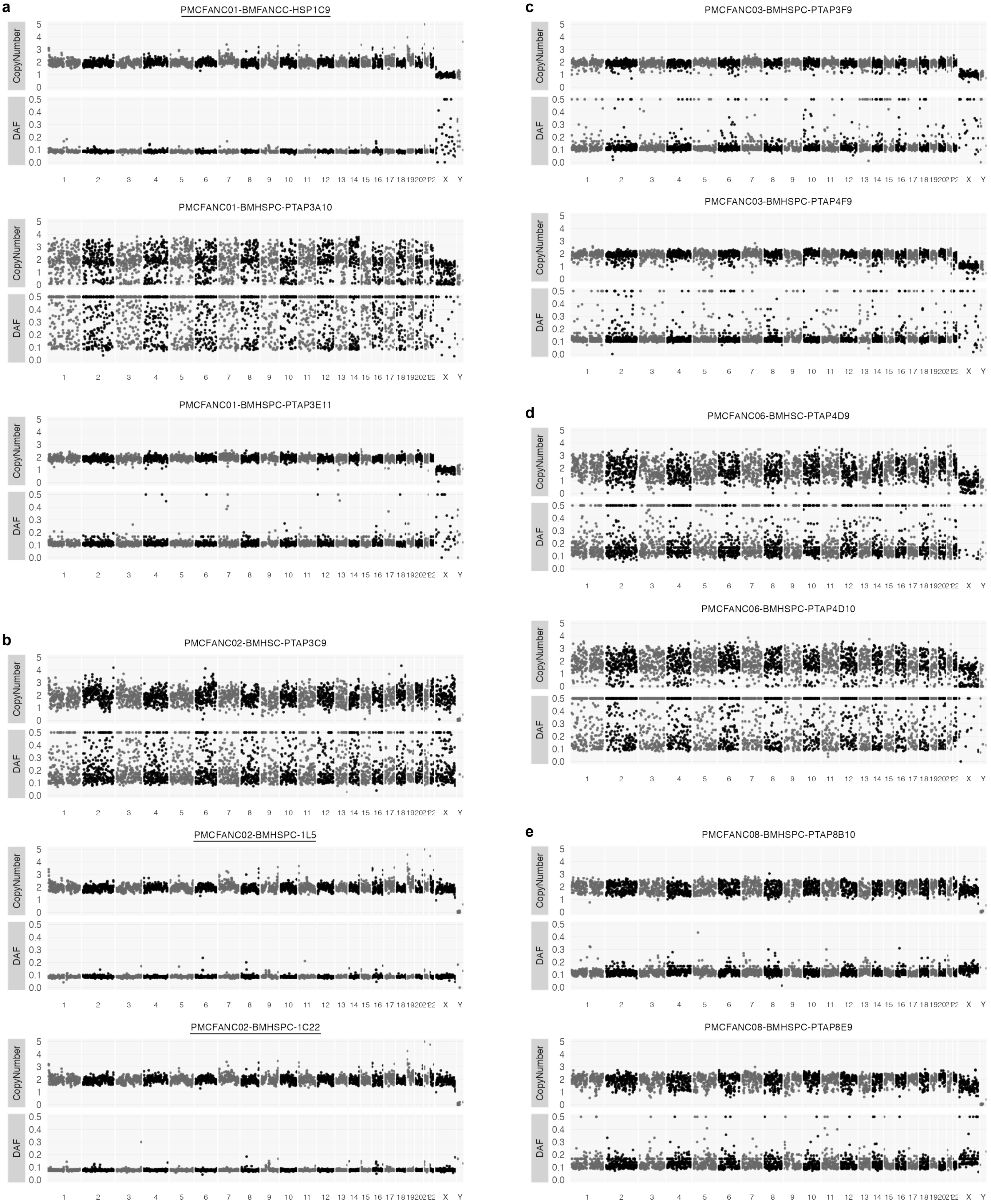


**Figure S9. No large chromosomal rearrangements detected in the HSPCs of patients with FA. a-e,** Copy number and DAF plots (at 1Mb resolution) after PTATO filtering of 12 analyzed HSPCs of 5 patients with FA. There is variability in the PTA quality between the single cells, leading to a lower sensitivity to detect SVs in some samples with low quality. Names of samples that were analyzed by WGS after clonal expansion (instead of PTA) are underlined.
